# Multiplexed dissection of a model human transcription factor binding site architecture

**DOI:** 10.1101/625434

**Authors:** Jessica E. Davis, Kimberly D. Insigne, Eric M. Jones, Quinn B Hastings, Sriram Kosuri

## Abstract

In eukaryotes, transcription factors orchestrate gene expression by binding to TF-Binding Sites (TFBSs) and localizing transcriptional co-regulators and RNA Polymerase II to cis-regulatory elements. The strength and regulation of transcription can be modulated by a variety of factors including TFBS composition, TFBS affinity and number, distance between TFBSs, distance of TFBSs to transcription start sites, and epigenetic modifications. We still lack a basic comprehension of how such variables shaping cis-regulatory architecture culminate in quantitative transcriptional responses. Here we explored how such factors determine the transcriptional activity of a model transcription factor, the c-AMP Response Element (CRE) binding protein. We measured expression driven by 4,602 synthetic regulatory elements in a massively parallel reporter assay (MPRA) exploring the impact of CRE number, affinity, distance to the promoter, and spacing between multiple CREs. We found the number and affinity of CREs within regulatory elements largely determines overall expression, and this relationship is shaped by the proximity of each CRE to the downstream promoter. In addition, while we observed expression periodicity as the CRE distance to the promoter varied, the spacing between multiple CREs altered this periodicity. Finally, we compare library expression between an episomal MPRA and a new, genomically-integrated MPRA in which a single synthetic regulatory element is present per cell at a defined locus. We observe that these largely recapitulate each other although weaker, non-canonical CREs exhibited greater activity in the genomic context.

## Introduction

The ability for organisms to precisely control gene expression levels and responses is crucial for almost all biological processes. Expression levels are controlled by *cis*-regulatory elements such as promoters and enhancers, *trans*-acting factors such as transcription factors (TFs), and cell, epigenetic and environmental states. Cis-regulatory elements help direct transcription responses by localizing and orchestrating interactions between active transcription factors, co-regulators, and RNA Polymerase II (ENCODE Project Consortium, 2012; Lambert et al., 2018). For control of human gene expression, a variety of large-scale projects seek to determine gene expression levels across various cell lines and cell types (Lizio et al., 2015, 2017), identifying functional elements that might control expression (ENCODE Project Consortium, 2012), the genome-wide characterization of epigenetic states of DNA (Roadmap Epigenomics Consortium et al., 2015), and the binding specificities of transcription factors (Jolma et al., 2013, 2015; Yin et al., 2017; Zhu et al., 2018). Collectively, while these efforts generally give us a parts list of putatively functional elements, understanding how these parts define quantitative levels of expression is still not well understood.

The combination of sequence motifs that recruit TFs, or TF-binding sites (TFBS), functionalize cis-regulatory elements via unique arrangements that help determine quantitative regulatory responses (Lambert et al., 2018; Spitz and Furlong, 2012). The consequences of subtle changes to TFBS compositions can be drastic. For example, clusters of weak-affinity Gal4 sites in yeast promoters increases expression synergistically, while stronger-affinity sites contributing to expression additively (Giniger and Ptashne, 1988). There can also be differences in TF occupancy of similar sequences in the genome that follow differences in the GC content of the surrounding sequence (Dror et al., 2015). Additionally, the placement of TFBSs can be highly conserved in close proximity to core transcriptional machinery, such as surrounding transcription start sites (TSSs) of genes (Tabach et al., 2007), and such placement can be critical for transcriptional activity (Kim and Maniatis, 1997; Kim et al., 1998). Lastly, the positional arrangement of TFBS combinations within cis-regulatory elements can modulate TF binding strength (Jolma et al., 2013, 2015) and TF activity can vary across the composition of TFBS combinations (Stampfel et al., 2015). Deciphering the logic imbued in cis-regulatory elements is difficult, as the limited set of natural variants and cell types are typically insufficient to control for variables such as sequence composition, TFBS composition and arrangements, and activity of trans-acting factors. Proving that particular sequences have causative effects on gene expression requires carefully controlled and high-throughput reverse-genetic studies.

The emergence of the massively parallel reporter assay (MPRA) allows for the testing of such reverse genetic transcriptional assays, and has become a powerful tool for the large-scale functional validation of regulatory elements across genomic and organismal contexts (White, 2015). These assays utilize the scale of synthetic DNA libraries and next-gen sequencing to determine the expression of thousands of individual regulatory elements in pooled expression measurements, enabling high-throughput functional characterizations of cis-regulatory logic. MPRAs have been used to quantify the transcriptional strengths of cis-regulatory elements and identify the motifs integral to element activity (Ernst et al., 2016; Kheradpour et al., 2013). Furthermore, several groups are using these systems to dissect how individual TFBSs drive quantitative regulatory responses in bacteria (Belliveau et al., 2018), yeast (van Dijk et al., 2017; Gertz et al., 2009; Levo et al., 2017; Sharon et al., 2012), human cell lines (Fiore and Cohen, 2016; Grossman et al., 2017; Weingarten-Gabbay et al., 2019), and animals (Kwasnieski et al., 2012; Smith et al., 2013; White et al., 2016). Collectively, these studies have begun to dissect TFBS logic by exploring how the regulatory grammar of different site combinations, numbers, and placements affect transcriptional activity.

Here we focus on how a range of factors guiding cis-regulatory architecture shape the activity of a single TFBS, the c-AMP Response Element (CRE). The CRE Binding (CREB) protein binds CRE and drives expression downstream of adenylyl cyclase activation (Gonzalez and Montminy, 1989; Montminy et al., 1986) across most cell types (Mayr and Montminy, 2001). CRE is ideally suited for exploring associations between TFBS architecture and regulation due to its ability to drive expression without other TFBSs in regulatory elements (Melnikov et al., 2012) and its ease of inducibility in a cell (Gonzalez and Montminy, 1989; Montminy et al., 1986), allowing finer control over active concentrations of the CREB protein. The most conserved, and likely to be functional, CREs generally localize within 200 basepairs (bp) of a TSS in the human genome (Mayr and Montminy, 2001; Zhang et al., 2005). Additionally, a previous MPRA that performed scanning mutagenesis on a commercial CRE reporter found mutations to CREs in closer proximity to the promoter had a greater effect on expression in addition to mutations to sequences flanking CREs (Melnikov et al., 2012). Here we explore the relationship between CRE’s distance to promoter elements and its activity in greater detail. We further explore the role other regulatory features play in modulating CREB protein activity including: CRE affinity, number, the spacing between multiple CREs, and the surrounding sequence content. Finally, although many MPRAs are performed episomally due to their ease and quickness (Fiore and Cohen, 2016; Grossman et al., 2017; Kheradpour et al., 2013; Melnikov et al., 2012), it’s been observed that episomal cis-regulatory element expression does not always correlate with their genomic counterparts (Inoue et al., 2017; Klein et al., 2019). Thus, we test our library both transiently and in a newly developed, singly-integrated genomic MPRA to better understand the quantitative and sometimes subtle differences between genomic and episomal assay context.

## Results

### CRE MPRA design and assay

We designed libraries with one or more CRE(s) by replacing sequence within three putatively inactive 150 bp background sequences to assay a range of features contributing to TFBS architecture (Figure 1A). These backgrounds were adapted from sequences with little reported activity from the Vista Enhancer Database (Visel et al., 2007) or a commercial reporter modified by removing previously identified CREs (Fan and Wood, 2007). We generated regulatory variants by replacing 12 bp regions of the backgrounds with either the consensus CRE (AT *TGACGTCA* GC), in which the central 8 bp region binds a CREB dimer (two monomer binding sites), or a weaker CRE (AT *TGA<u>A</u>GTCA* GC), where one of the central dinucleotides bound by both monomers was mutated and has been previously shown to reduce activity (Mayr and Montminy, 2001; Melnikov et al., 2012). For the majority of analysis, we used two CRE libraries. The first library, the CRE Spacing and Distance Library, assays CREB activity as a function of both the spacing between CREs and CRE distance to the minimal promoter by moving two consensus CREs across the 150 bp backgrounds at six defined spacings (0, 5, 10, 15, 20 and 70 bp) between the two sites. In the CRE Number and Affinity Library, we explore the effect of both CRE number and affinity upon expression by designating 6 equally-spaced locations across all backgrounds in which each location is replaced with either the weak CRE, consensus CRE, or no CRE.

**Figure 1.**
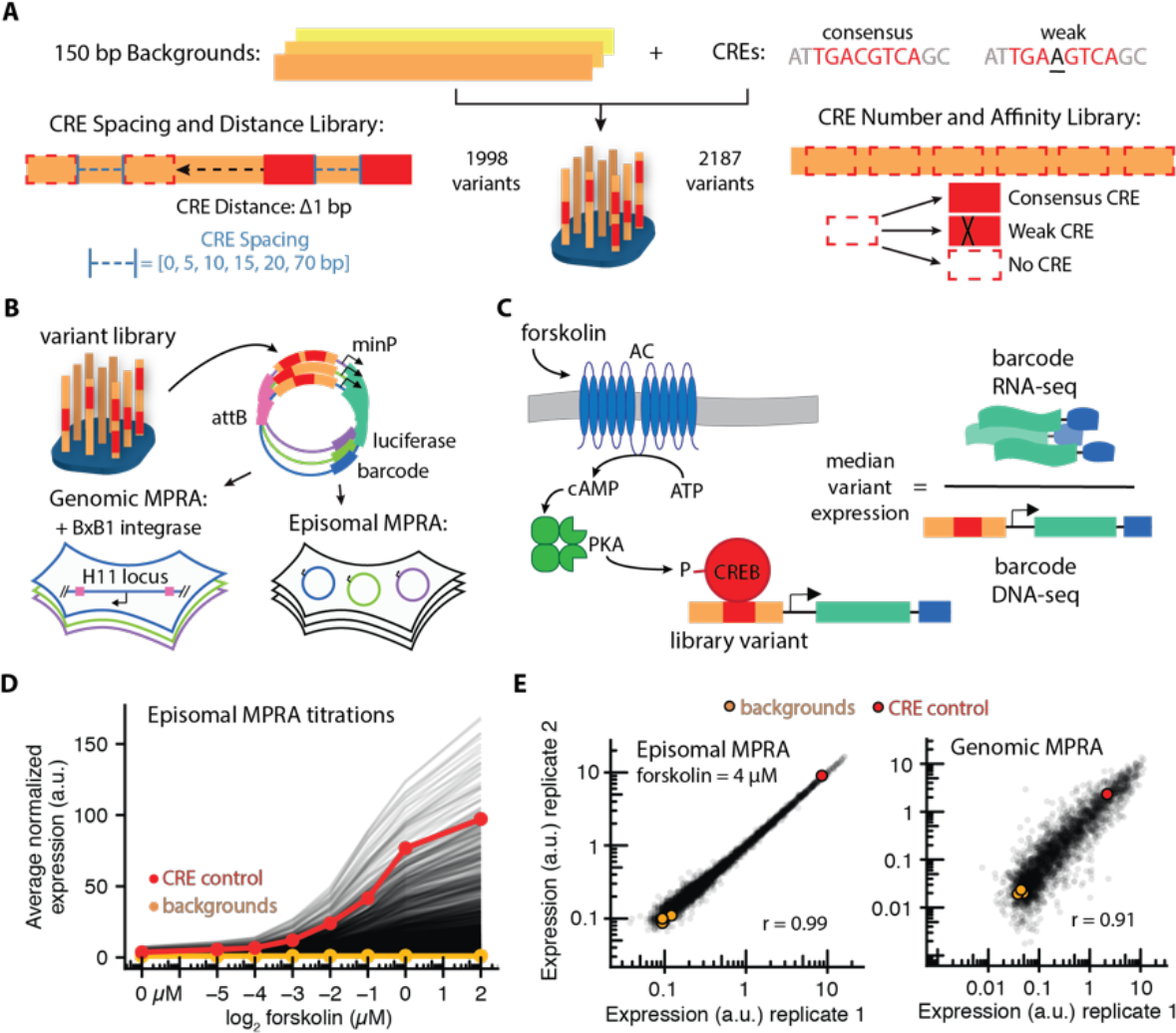
CRE regulatory library design, synthesis, and assays. (A) We replaced sequence within three putatively inactive backgrounds with consensus and/or mutant CREs to generate two libraries with varying spacing and distance to the minimal promoter (CRE Spacing and Distance Library) or the number and strength of the binding (CRE Number and Affinity Library). (B) We assembled the library reporter vectors with variants upstream of a minimal promoter, luciferase ORF, and a unique barcode sequence. The pool of library reporter vectors was assayed in a HEK293T cell line either by transient transfection or integrated at one allele of the Human H11 locus. (C) We stimulated CREB protein activity with forskolin, which activates cAMP signaling. Expression levels were determined as the ratio of barcodes reads in the RNA to that of the DNA sample. (D) Library expression following increasing forskolin concentrations in the episomal MPRA. Each variant is normalized to the expression of their corresponding background (no CREs), then averaged across replicates. (E) Variant expression exhibits strong correlation between biological replicates in both MPRAs, here shown at maximally-inducing concentrations of forskolin in both assays (episomal r = 0.99 and genomic r = 0.91).

We used Agilent OLS synthesis to construct our designed libraries, added random 20 nt barcodes to the 3’ end, and mapped these barcode-variant associations. For MPRA analysis we only considered barcodes corresponding to perfect matches to our designs, identified at this stage via next-gen sequencing. We cloned these libraries into a reporter construct we engineered to maximize signal to noise when integrated into the genome (Supplemental Fig. 1C) and then cloned a minimal promoter and luciferase gene between variant and barcode, placing the barcode in the 3’ UTR of the luciferase gene. The assays were conducted at varying induction conditions, either episomally by transient transfection (Episomal MPRA), or singly-integrated into the intergenic H11 safe-harbor locus (Zhu et al., 2014) using BxBI-mediated recombination (Genomic MPRA) (Duportet et al., 2014; Jones et al., 2019; Matreyek et al., 2017; Xu et al., 2013) (Figure 1B). The episomal MPRAs were run in biological duplicate across 14 different concentrations of forskolin (Figure 1D, Supplemental Fig. 1E, and Supplemental Fig. 2), which stimulates phosphorylation and activation of the CREB protein by activating adenylyl cyclase (Gonzalez and Montminy, 1989). The genomic MPRA was run in biological duplicate at full induction (Supplemental Fig. 1D). After forksolin stimulations, we isolated barcoded transcripts from cells and used next-gen sequencing to determine barcode prevalence per RNA samples and plasmid (episomal MPRA) or genomic (genomic MPRA) DNA samples. Since each variant was mapped to multiple barcodes, we first determined the expression of each barcode via the ratio of normalized reads in the RNA over the DNA sample. We then determined variant expression from the median expression of all barcodes mapped to each variant. Both episomal and genomic MPRAs indicated high reproducibility between separately stimulated replicates (Figure 1E, episomal Pearson’s *r* = 0.99, genomic *r* = 0.91, and Supplemental Fig. 1E). In both assays, the difference between backgrounds alone and a positive CRE control adapted from a commercially-available reporter plasmid (Fan and Wood, 2007) spanned the majority of expression variation amongst variants.

### The role of CRE spacing and distance on expression

We explored the extent to which positioning of CRE within a regulatory element quantitatively affects its transcriptional activity. We initially assayed the relationship between CRE distance relative to a downstream promoter and variant expression using 1 consensus CRE in a separate library, but found it drove minimal expression in the episomal MPRA after CREB activation (Supplemental Fig. 3). We then examined the expression driven by the CRE Spacing and Distance Library. This library varied the relative positioning of two consensus CREs with respect to the minimal promoter (referred to as CRE *distance*), and altered the number of nucleotides between the two sites (referred to as CRE *spacing*) (Figure 2A). We tested *spacings* of 0, 5, 10, 15, 20 or 70 bp between the two CREs, and then tested *distance* by moving these sites with each of the *spacings* across the backgrounds one base at a time, spanning the 150 bp backgrounds. We see activation occurs in a dose-dependent manner in the episomal MPRA, with a ~10 bp expression periodicity which was more apparent at higher concentrations of forskolin. This 10 bp periodicity was consistent across CRE *spacings* and backgrounds, displayed similar patterns between the genomic and episomal assays, but differed between backgrounds (Supplemental Fig. 4). Such periodicity has been observed before for single TFBSs in a variety of model systems (Kim et al., 2013; Sharon et al., 2012; Takahashi et al., 1986). In addition, we also observed a general decrease in expression as CRE *distance* increased. Across backgrounds, MPRA formats, and CRE *spacings*, the change in median expression between the CRE *distance* range of 67-96 bp and 147-176 bp resulted in a median 1.5 to 2.2-fold decrease in expression, with larger effects observed in the genomic MPRA (Figure 2B).

**Figure 2.**
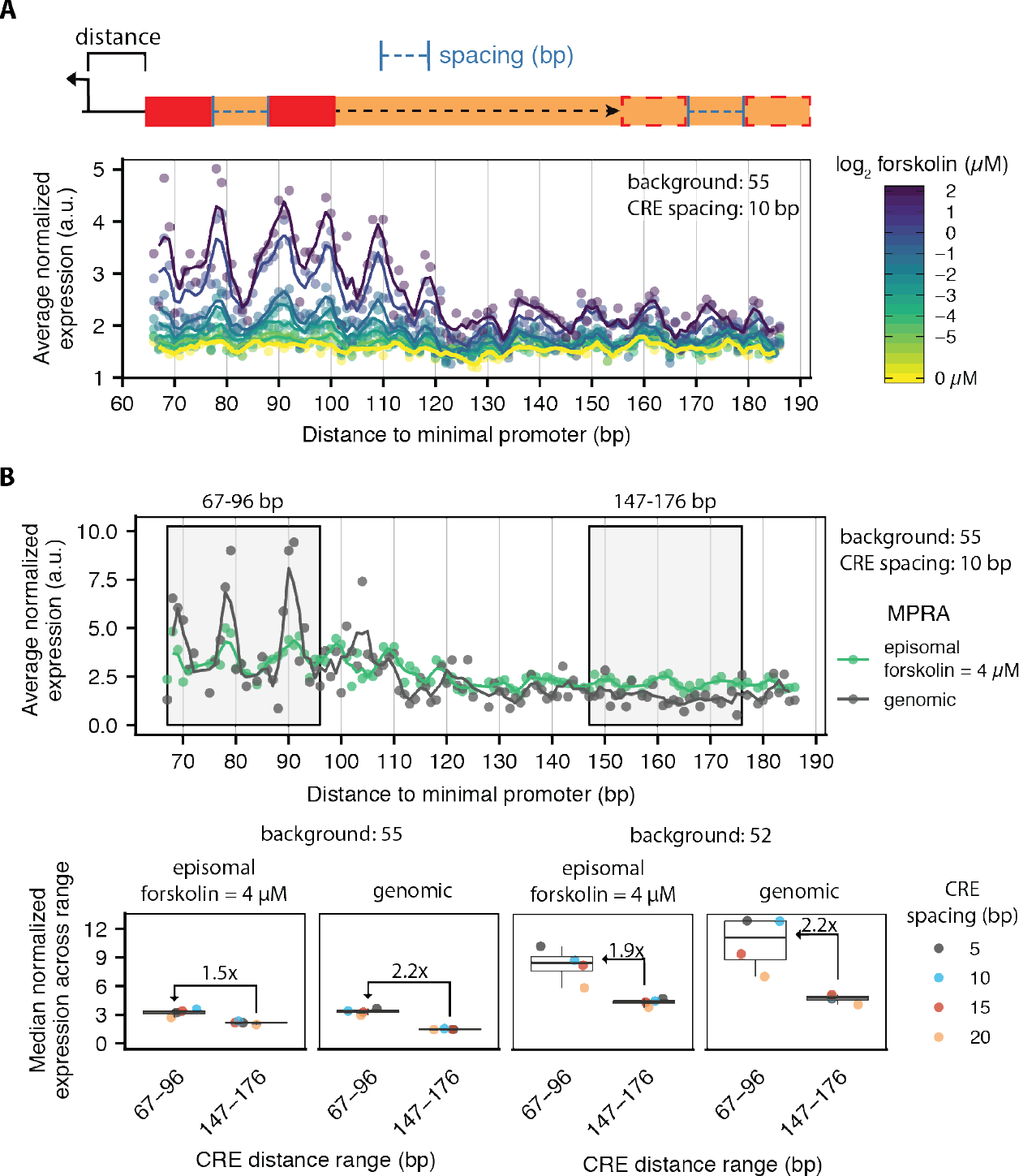
CRE proximity to promoter elements is associated with higher expression. (A) The expression profiles for two CREs that are 10 bp apart display a periodic signal as they are moved away from the minimal promoter for Background 55 (with forskolin concentration shown in color). The lines are 3 bp moving averages of the points. (B) Expression profiles for variants with Background 55 and 10 bp CRE *spacing* at maximally-inducing concentrations for both episomal and transient MPRAs (top panel). Expression decreases (lower panel) as the distance of the CREs from the proximal promoters increases across the backgrounds, spacings, and MPRA formats (as measured by the median expression across *distance* ranges 67-96 bp and 147-176 bp).

We noticed an offset in expression periodicity according to varied CRE *spacing* that was most pronounced in variants with background 41 in the episomal MPRA (Figure 3A). In particular, 5 and 15 bp CRE *spacings* exhibited similar local expression maxima at CRE *distances* 79 and 88 bp, whereas 10 and 20 bp *spacings* had maxima at 83 and 92 bp. This ~5 bp shift was also observed in background 55, albeit at different *distances* (Supplemental Fig. 5); it is of note we did not observe similar periodicity alignments and offsets in a genomic context. In these instances, we indicate *distance* from the minimal promoter to the start of the first CRE, such that the proximal CRE is in the same position across all expression profiles. Thus, the only differences between variants that may be causing this periodicity shift is the altered placement of the distal CRE following CRE *spacings*. Using the CREB bZIP structure bound to CRE (Schumacher et al., 2000), we modeled 2 dimers bound to CREs with 5 and 10 bp *spacings* (discussed in Methods) by aligning protein-DNA density to DNA backbone (Figure 3B). This simplified model does not incorporate density from full-length CREB proteins or both proteins’ effects on local DNA bending. A 5 bp shift in *distances* driving expression maxima between 5 and 10 bp *spacings* corresponds to about half a helical turn of B-form DNA (10.4 bp/turn). Our model positions the proximal CREB dimer (1) on the opposite face of the DNA helix between 5 and 10 bp *spacings* when they are both at their expression maxima. On the other hand, the distal CREB dimer (2) would be similarly oriented between the 5 and 10 bp *spacings*, but at a full helical turn distance from one another on the DNA.

**Figure 3.**
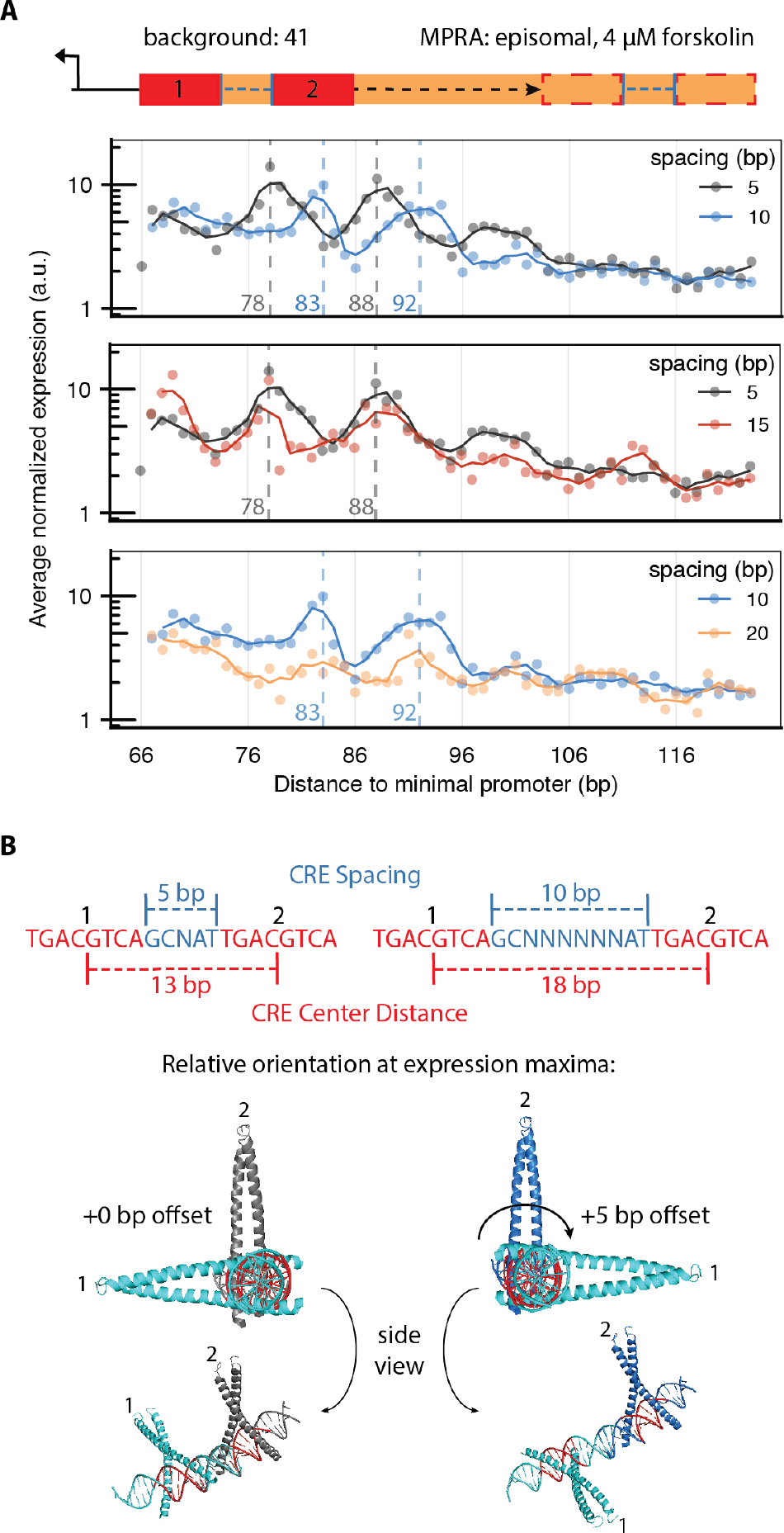
CRE *spacing* modulates expression periodicity as CRE *distance* is varied. (A) Average background-normalized expression of variants with background 41 in the episomal MPRA plotted as a function of CRE *distance*. Solid lines correspond to the 3bp moving average estimate and dashed lines indicate local expression maxima determined from 5 and 10 bp CRE *spacings*. Overlays indicate the offset of expression periodicities between 5 and 10 bp *spacings* and alignment between both 5 and 15 and 10 and 20 bp *spacings*. (B) Using the published structure of a CREB∷bZIP dimer bound to a CRE (PDB: 1DH3), we modeled the expected positioning of CREB dimers bound to the CRE proximal (1) and distal (2) to the promoter for both the 5 and 10 bp CRE *spacings* at the CRE *distance* of their respective local maximum. This model approximates the distal dimer (2) in similar orientations at the two local expression maxima, with the proximal dimer on opposite faces of the DNA.

### The role of CRE number and affinity upon expression

While CRE *distance* and *spacing* in regulatory elements help shape CRE’s activity, the overall number and affinity of CREs likely plays a larger role in determining expression. The design of the CRE Number and Affinity Library assays these effects while taking into consideration the contribution of CRE position. Each variant in this library contained unique combinations of consensus and weaker-affinity CREs spanning an assortment of 6 positions along the backgrounds (Figure 1A). A constant 17 bp CRE *spacing* was implemented in this library design in order to sample a range of predicted CREB protein orientations along the DNA across the 6 positions (Supplemental Fig. 6). Even so, the number of consensus CREs alone largely determined variant expression and this relationship followed a non-linear increase and eventual plateauing of expression in both MPRA formats (Figure 4A and Supplemental Figs. 7–8). We observed a similar increase with the number of weak CREs if at least one consensus CRE was also present within the variant, although this effect varied per background and between episomal and genomic MPRAs. While the number of consensus CREs largely determined variant expression, there was a large amount of expression variability per arrangement of similar numbers of consensus and weak CREs, perhaps due to combinations of CREB protein orientations. To explore how the different arrangements of CREs across the six positions shaped expression per CRE combination, we fit a log-linear model of expression to the independent contributions of CREs at each position (Figure 4B). We allowed different weights to be fit per CRE affinity per position and also included an independent background term to account for expression differences between backgrounds.

**Figure 4.**
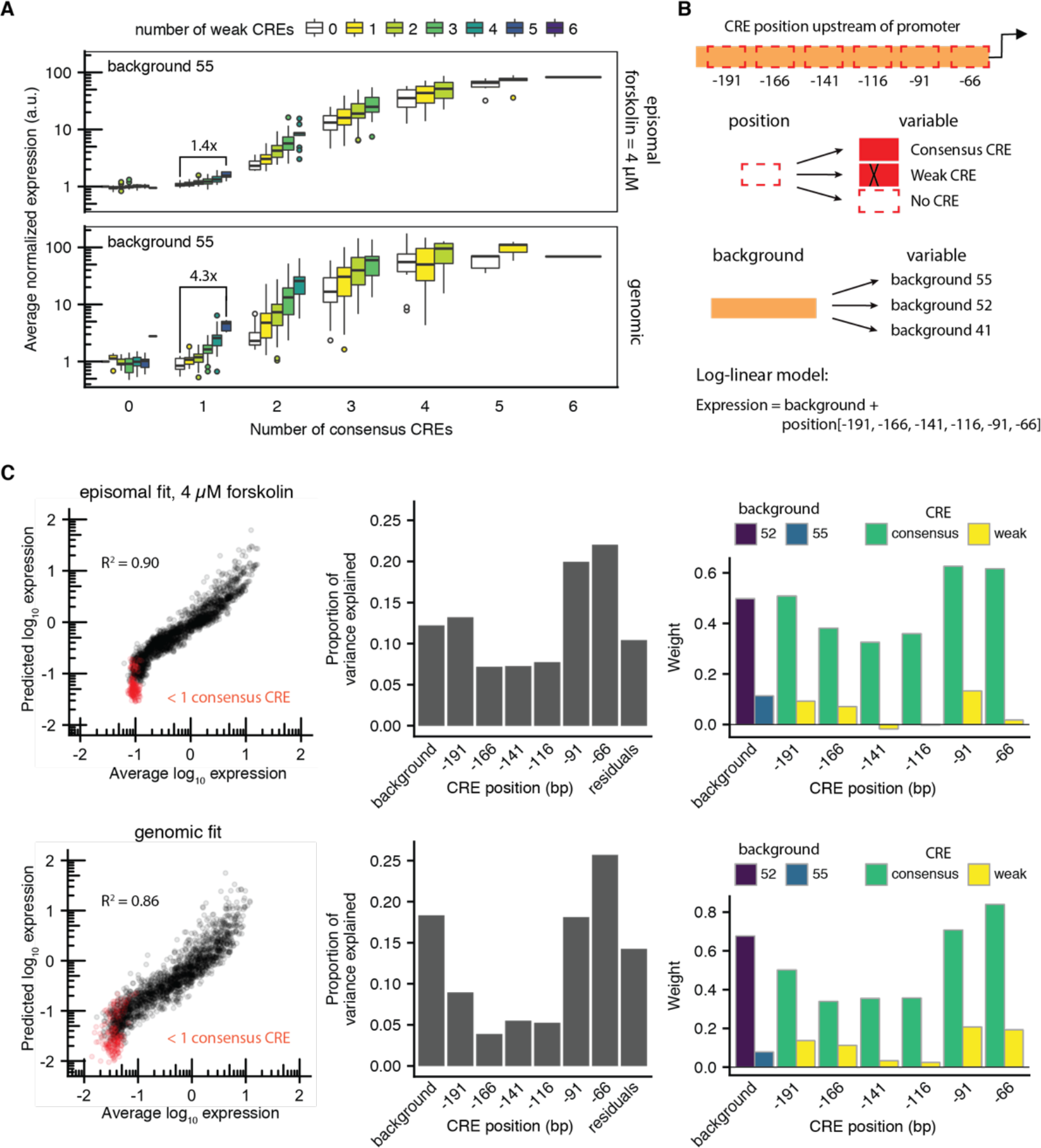
CRE number and affinity largely determines variant expression with variation explained by CRE position and background. (A) The expression of variants with background 55 according to their number of consensus (x-axis) and weak (colored subsets) CREs in the integrated and episomal MPRAs. The change in median expression between variants with 1 consensus CRE and those with 1 consensus and 5 weak CREs is indicated. (B) A simple linear model was fit to log-transformed expression using the identities of the background and TFBSs at each position as inputs. (C) This fit model correlates well with measured expression for episomal (*R*^*2*^ = 0.90) and genomic (*R*^*2*^ = 0.86) MPRAs and deviates from measurements of variants with no consensus CREs (left panel). Analysis of variance indicates 90% episomal and 86% genomic variance in expression is explained by the model. CREs occupying the two closest positions to the promoter had the strongest effects. The weights of categorical variables show the relative effects of strong and weak CREs and the effect of each background relative to variants with no CREs and with background 41.

We found this independent, position-specific model explained a majority of expression (Figure 4C, left panels) in both the episomal (r = 0.95) and genomic MPRAs (r = 0.93). Although the model was inaccurate at predicting activity from low-expressing variants, this was largely due to variants with weak CREs and no consensus CREs driving little variation in activity in our assay. Accordingly, we found that CREs closest to the promoter (66 and 91 bp upstream of the promoter) explained 42.6% of the variance in the episomal MPRA and 43.8% in the genomic MPRA (Figure 4C). This is expected as both positions fell within the ~110 bp of higher expression observed with the CRE Spacing and Distance library. None of the weights fit to CRE positions followed a trend in predicted CREB protein orientations (Supplemental Fig. 6), thus CRE’s distance to promoter elements may mask the subtle effects of CREB protein orientation previously observed with 2 CREs. Background alone explained 12.1% of the variance episomally and 18.4% of genomic expression variance. The trend in weights fits per background did not follow overall GC content, nor the total CG dinucleotides (not shown), but may follow GC content similarity to that of the CRE itself (50% GC). Overall, while we find that the number of consensus CREs per variant largely determined expression, the combination of CRE positions along the backgrounds and the backgrounds themselves explained a majority of expression in our assays.

### Differences between episomal and genomic MPRAs

There were a number of differences between the genomic and episomal expression trends resulting from increasing CRE numbers per variant. First, variants with weak CREs drove greater expression in the genomic MPRA, especially within the context of few consensus CREs per variant. For example, the change in median expression between variants with one consensus CRE and those with one consensus CRE and five weak CREs within background 55 was 1.4-fold in the episomal MPRA and 4.3-fold in the genomic MPRA (Figure 4A). Second, expression plateaued at about six consensus CREs for most backgrounds in the episomal MPRA, while in the genomic MPRA, this occurred at about four consensus CREs for all backgrounds (Figure 4A and Supplemental Fig. 8). To explore differences in expression trends between the two MPRAs, we compared expression between the two assays according to different combinations of CRE affinities within variants (Figure 5A). There was a strong linear relationship between genomic and episomal expression for variants containing one to six consensus CREs (Figure 5A left panel, *R*^*2*^ = *0.92*). Using this linear relationship as a reference (red line), we then compared expression between the two assays for variants containing one to six weak CREs (Figure 5B middle panel) and combinations of one to six consensus and weak CREs (Figure 5B right panel). Expression deviated from this linear relationship for variants containing weak CREs, with 79% of the variants exhibiting higher relative expression in the genomic MPRA.

**Figure 5.**
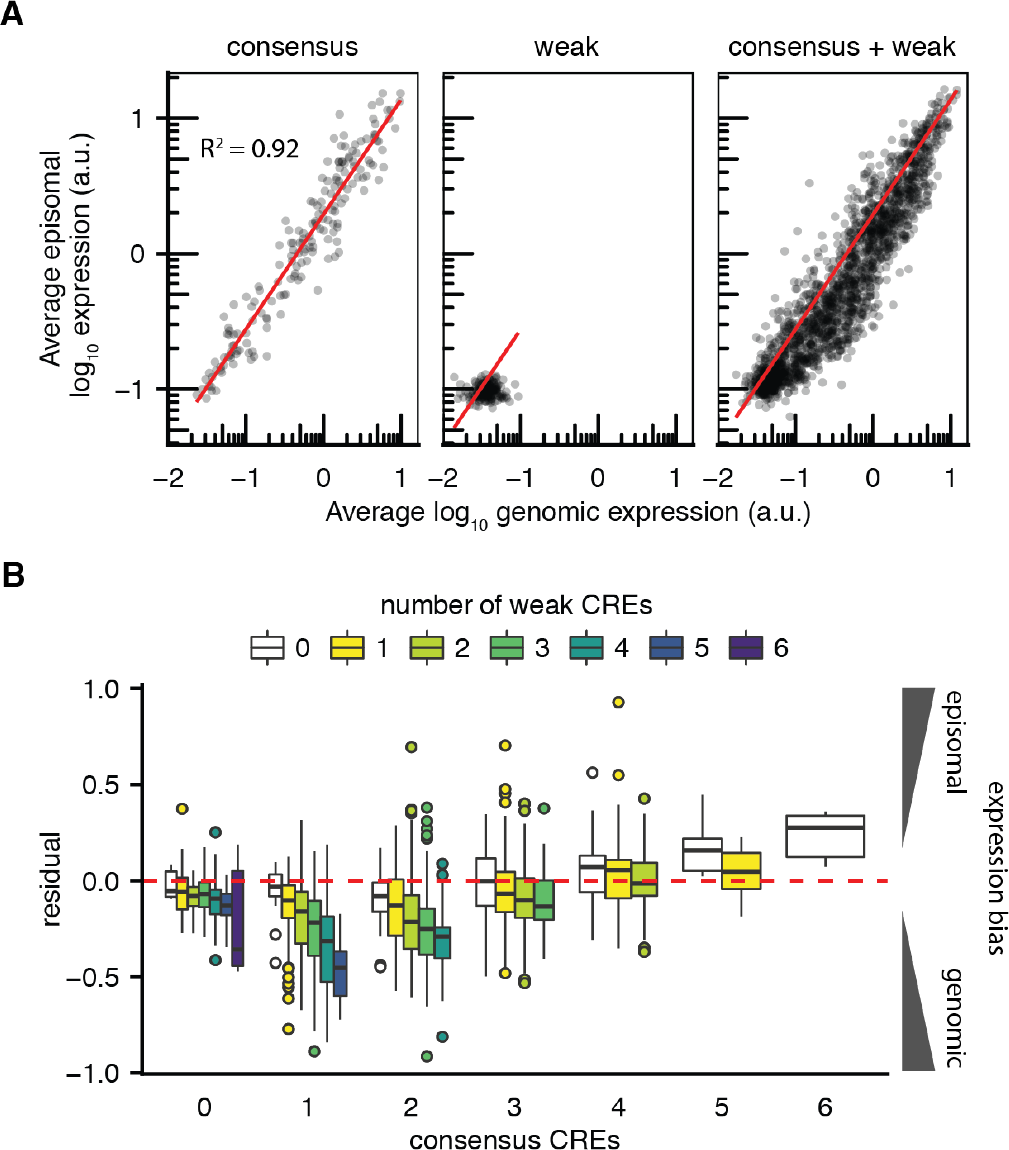
Variants with weak CREs exhibit higher relative expression in a genomic context. (A) Variants across all backgrounds subset according to site affinity composition. Subsets include variants with 1-6 consensus CREs (left panel), 1-6 weak CREs (middle) and combinations of 1-6 weak and consensus CREs (right). Variants with only consensus CREs drive similar relative expression (*R^2^ = 0.92*, red line) between genomic and episomal MPRAs. Most variants with weak CREs drive higher relative expression in the genomic MPRA compared to variants without weak CREs. (B) Variant expression according to their number of both consensus (x-axis) and weak CREs (colored subsets). Using the expression correlation line between variants with 1-6 consensus CREs in both MPRAs (red line) as a reference, the residual of each variant to this line is plotted along the y-axis. Variants containing higher numbers of weak CREs drive higher relative expression in the genomic MPRA while those with higher numbers of consensus CREs drive higher relative expression in the episomal MPRA.

When we broke down this comparison according to numbers of consensus and weak CREs per variant, we observed higher relative genomic expression of weak CREs in the context of few (0-3) consensus CREs and higher relative episomal expression with combinations of at least 4 consensus CREs (Fig. 5B). This effect may be due to the differences in CRE variant copy numbers between assays. Lipofectamine transfections can result in 10^1^-0^4^ plasmids/nucleus (Cohen et al., 2009). In this context, high numbers of variants with many consensus CREs may out-compete variants with weak CREs for CREB-protein binding. In similar systems, the titration of plasmids containing the same TFBS as a single chromosomal reporter has altered the expression of the chromosomal reporter (Lee and Maheshri, 2012) in a manner dependent upon the strength of the plasmid “competitor” sites and cellular TF concentrations (Brewster et al., 2014). Although multi-copy MPRAs are ideal for assaying large libraries of variants by reducing the number of cells required to cover library diversity, expression may be interpreted in the context of variant competition when variants contain similar TFBSs. Despite this, we show MPRAs that assay many variants per cell can largely recapitulate other TFBS features shaping genomic expression driven from single variants.

## Discussion

By performing synthetic manipulations of a single TFBS, the CRE, we present here a characterization of various regulatory rules governing TF activity. In particular, we assay the effects of CRE number, affinity, distance to promoter, and the spacing between multiple CREs within regulatory elements. Additionally, we show how a subset of these features shape expression when tested in combination. The limited complexity of natural cis-regulatory elements would not have allowed us to explore these features at bp-resolution and in such a controlled manner. Thus, we chose to isolate these features using synthesized regulatory elements in the format of an MPRA. Furthermore, we integrated variants at a single copy per cell within the same genomic environment to better approximate genomic expression. We present here improvements in a single-copy, defined-locus genomic MPRA performed in a human cell line, indicating the feasibility of reproducible expression measurements of lowly-expressing transcripts.

Although the variants assayed here are synthetic, we expect the regulatory trends observed to define expression from natural cis-regulatory elements as well. We observed a drop in transcription with placements of CRE beyond 120 bp *distance* to our minimal promoter, following similar findings with manipulation of native CREs (Tinti et al., 1997). Yet diminished expression periodicity was still observed at *distances* up to ~180 bp (Figure 2 and Supplemental Fig. 4). CREB protein recruits RNA polymerase II through interactions with TFIID, while it’s phosphorylated form drives polymerase isomerization and transcription (Kim et al., 2000). Thus CRE’s proximity to promoter elements may be integral to polymerase recruitment and transcription, perhaps explaining CRE’s enriched localization within 200 bp of TSSs in the human genome (Mayr and Montminy, 2001; Zhang et al., 2005). TF’s that also directly recruit and activate RNA Pol II may also follow similar trends in activity according to TFBS promoter proximity.

In addition to overall distance effects, CRE’s precise positioning likely plays a role in its activity in natural regulatory elements via the periodicity observed. In line with CRE’s localization around TSSs, conserved TFBSs that exhibit location-specificity in the genome are mostly found between 200 bp upstream and 100 bp downstream of a TSS (Tabach et al., 2007). For instance, manipulations that place the highly conserved 8 TFBSs in the INF-ß enhanceosome at half a helical turn from the original 47 bp upstream of the TSS (Panne, 2008) reduce transcription (Kim and Maniatis, 1997). These altered enhanceosome orientations hinder TF recruitment of TFIIB and coactivator CREB protein binding protein (CBP), which binds RNA Pol II (Kim et al., 1998). While CREB protein similarly drives transcription through interactions with CBP (Zhang et al., 2005), it is unclear how such interactions would drive the periodicity offset observed with 2 CREs in local proximity. In another study, a cyclical relationship was observed between expression and the varied spacing of 2 ZEBRA bZIPs upstream of a TATA element, with minimal activity observed when both dimers were simulated to be in-phase (Huang et al., 2012). Thus, we may be observing the manifestation of this pattern when both CREs with fixed spacing are placed at varying distances away from the TSS. While predicted CREB protein orientations did not seem to influence expression across combinations of up to 6 CREs, it nevertheless may still play a role if this library was tested across varied CRE *distances*.

The number of consensus CREs largely determined expression in our assays, following similar trends as other homotypic clusters of TFBSs assayed (van Dijk et al., 2017; Gertz et al., 2009; Sharon et al., 2012; Weingarten-Gabbay et al., 2019). In contrast, a recently published MPRA in human cell lines found increasing number of CREB binding sites did not increase expression, although these sites were assayed in the absence of forskolin (Weingarten-Gabbay et al., 2019). Although the number of sites generally increase expression in both MPRAs tested here, there are instances in which variants drive less expression following increasing consensus CRE number. In the genomic MPRA for example, many variants of a particular background drive higher expression with 4 consensus CREs than that of variants with 5 or 6 (Figure 4A and Supplemental Fig. 8). In some of these examples, higher expression is observed with the addition of a weak CRE to a variant as opposed to a consensus CRE. Closely-spaced TFBSs can restrict the diffusion of TFs along DNA if one is already bound (Hammar et al., 2012), leading to binding competition between binding sites. This competition has been implicated in decreasing expression in a similar TFBS MPRA (van Dijk et al., 2017) and may explain our observations in the genomic MPRA. This effect is not as apparent in the episomal MPRA, a feature that may be explained by predominantly measuring competition between plasmids over competition between CREs in a single variant.

Lastly, we provide further evidence MPRA design and regulatory context must be considered in characterizations of regulatory features shaping expression. In both MPRAs, the contribution of background to variant expression is more predictive of variant activity than the presence of CRE at many positions along these backgrounds. The surrounding sequence content may play a similarly significant role for many other TFBSs, especially those that exhibit a bias in binding events based on the GC content similarity of the surrounding sequence to that of the TFBS itself (Dror et al., 2015). Therefore, we recommend incorporating multiple sequences as backgrounds in similar synthetic regulatory element designs especially since the use of a single background, as has been employed in many MPRAs, can influence TFBS trends observed (Supplemental Fig. 8). Additionally, we indicate here the ability of episomal assays to approximate genomic regulatory rules, yet also warn of the potential pitfalls of transient assays. Overall, we observe strong correlation between our episomal and genomic MPRAs (Supplemental Fig. 9, Pearson’s r = 0.91). Yet the activity of variants with weaker-affinity CREs varies considerably between assays, which may be explained by differences in variant, and hence CRE, copy numbers in a cell. Thus we would not expect this effect to skew expression measurements of libraries assaying a high diversity of TFBSs. Alternatively, this could also be attributed to more consistent genomic structure in chromatin.

While we characterize various regulatory rules shaping the activity of a single TFBS, the CRE, we use this as a model to estimate a small fraction of the complexity of expression attained by combinations of TFBSs in natural cis-regulatory elements. Exploring how these rules scale with other TFBSs is integral to our understanding of cis-regulatory logic. Similar high-throughput approaches can build from the constraints explored here to develop more complex dissections of TFBS architectures. Transcriptional activation is thought to occur via phase-separated TF-coactivator-Pol II hubs, with local concentrations of these factors driving expression non-specifically (Boehning et al., 2018; Chong et al., 2018; Reiter et al., 2017). The interplay between transcriptional activity in these phase-separated systems and TFBS grammars needs further exploration. Further characterizations using similar synthetic systems will further our comprehension of cis-regulatory elements and our ability to confidently compose new ones with predictable activities.

## Acknowledgements

We thank the Kosuri Lab for providing useful feedback on earlier versions of this manuscript. We thank the UCLA Eli & Edythe Broad Center of Regenerative Medicine & Stem Cell Research Sequencing Cores and the Technology Center for Genomics and Bioinformatics for providing next-generation sequencing, as well as the UCLA Eli & Edythe Broad Center of Regenerative Medicine Flow Cytometry Core for assistance in flow cytometry. We also thank Laura Day from the Kruglyak lab at UCLA for use of and assistance on their MiSeq. We also thank Dr. Michael R. Sawaya for his guidance in CREB-CRE structural modeling.

## Data and materials availability

All custom scripts and code for figure generation are available at the Kosuri Lab Github and can be accessed at the following link: https://github.com/KosuriLab/CRE_library_code. Raw data is available, and will be made available after peer-review. Plasmids and cell lines are available upon request.

## Author Contributions

J.E.D. and S.K. conceptualized the experiments. K.D.I. computationally designed the oligo library. J.E.D. performed all experiments and most computational analysis with the help of K.D.I. E.M.J. helped with the generation and validation of the H11 landing pad integration vector and cell line generation and validation. Q.H. helped with library generation for Supplemental Figure 3. S.K. and J.E.D. wrote the manuscript with input from all authors.

## Materials and Methods

### CRE regulatory library design

The CRE regulatory library was designed using three 150 bp *backgrounds* as templates and either a consensus CRE, taken from the CREB1 sequence logo in the JASPAR database, or weaker-affinity CRE, in which one of the central dinucleotides important for binding of both CREB protein monomers was mutated (Mayr and Montminy, 2001; Melnikov et al., 2012). Two of the backgrounds were adapted from previous MPRAs (background 55 (Melnikov et al., 2012) and background 41 (Smith et al., 2013)) and a third was isolated from a human genomic region indicating minimal activity in the developing eye in the VISTA enhancer database (Visel et al., 2007) (background 52). Both background 41 and 52 were obtained from the human genome, with 41 corresponding to Chr9: 81,097,684-81,097,833 and 52 to Chr5: 89,377,854-89,378,003 from GRCh38. Background 55 corresponds to the CRE response element of a commercial reporter plasmid (Fan and Wood, 2007) with a portion duplicated to reach 150 nt and with all CREs scrambled, maintaining their GC content. Variants were generated by replacing background sequence with CREs along with a constant 2 bp flanking nucleotides to ameliorate local sequence effects due to CRE placement in the backgrounds (Levo et al., 2015). A MluI restriction enzyme site (ACGCGT) was placed upstream of each variant and KpnI restriction enzyme site (GGTACC) was placed downstream, each for library cloning. Lastly, a pair of 19 nt amplification primers (Eroshenko et al., 2012) specific to each of the 2 library designs were added to each design producing 200 nucleotide libraries of the format: (5’ −> 3’) subpool primer 1 - MluI - variant - KpnI - subpool primer 2. The resulting 4185 variants corresponding to the 2 designs in Figure 1, 417 variants corresponding to the single-CRE library, and one CRE positive control (Fan and Wood, 2007) were all synthesized on Agilent Microarrays.

### Library cloning

OLS libraries (Agilent Technologies, Santa Clara, CA) were resuspended to a final volume of 200 nM in TE pH 8.0 (Sigma-Aldrich, Saint Louis, MO). Libraries were amplified using 1 μL of a 10-fold dilution of the library, the respective subpool primer pairs (Subpool_#_F and Subpool_#_R with # representing the sub-libraries present, 2 refers to the single CRE Distance library, 3 refers to the CRE Spacing and Distance library, and 5 refers to the CRE Number and Affinity library) and with KAPA HiFi HotStart Real-time PCR Master Mix (2X) (Kapa Biosystems, Wilmington, MA) following the recommended cycling protocol at 14 cycles. Random barcodes were added to variants in a second PCR in which the primer downstream of variants contained 20 nucleotides of random sequence, synthesized with the machine-mixed setting (Integrated DNA Technologies, Coralville, IA). One ng product was used in this barcoding qPCR using biotinylated primers (SP#_Biotin and SP#_Biotin_BC_R with subpool numbers corresponding to library designs as before). The qPCR was performed with KAPA HiFi HotStart Real-time PCR Master Mix (2X) (Kapa Biosystems) for 11 cycles following the recommended cycling protocol. Barcoded libraries were digested with MluI-HF (New England Biolabs, Ipswich, MA) and SpeI-HF (New England Biolabs) in 1X cut-smart buffer (New England Biolabs). The biotinylated primers and undigested library members were removed using Dynabeads M-270 Streptavidin (Thermo Fisher Scientific, Waltham, MA), using the recommended “Immobilize nucleic acids” protocol, collecting the supernatant after adding the library mixture to the beads.

The barcoded and digested library members were cloned into the integration vector (pJDrcEPP) that had previously been digested using MluI-HF and SpeI-HF in 1X cut-smart buffer (New England Biolabs). Ligation was performed at a 1:3 ratio of pJDrcEPP:library using T4 DNA ligase (New England Biolabs). Ligation product was cleaned-up using a Clean and Concentrator Kit (Zymo Research, Irvine, CA) followed by drop-dialysis for 15 minutes with UltraPure™ DNase/RNase-Free Distilled Water (Thermo Fisher Scientific) before transforming 1 μL into NEB 5-alpha Electrocompetent E. coli (New England Biolabs) following the recommended protocol. Dilutions of transformants were plated on 50 μg/mL Kanamycin (VWR, Radnor, PA) LB plates at 10-fold dilutions to 1/10,000 and grown overnight at 37°C and the remainder of the cells were left in SOC (New England Biolabs) at 4°C overnight. The next day, dilutions were counted, estimating 695,000 and 950,000 original transformants for the placement and spacing library and the number and affinity library, respectively. The remainder transformants kept overnight at 4°C were pelleted and placed in fresh LB for 3 hours at 30°C, then diluted into 100x volume LB + 50 μg/mL Kanamycin (VWR) and grown at 30°C for 18 hours before isolating library vectors using QIAprep (Qiagen, Hilden, Germany) spin miniprep kits.

Library vectors (pJDrcEPP_lib) were then digested in 2 steps, in order to isolate plasmids correctly cut at the synthesized KpnI recognition site. Vectors were first digested with KpnI-HF along with rSAP (New England Biolabs) in 1X CutSmart buffer. Products were run on a 0.8% TAE agarose gel and linearized plasmids were isolated with Zymoclean Gel DNA Recovery Kit (Zymo Research). Vectors were then digested in a similar fashion with XbaI (New England Biolabs) without gel isolation. The minimal promoter and luciferase insert was prepared using biotinylated PCR primers (Amp_minPLuc2_Biotin_For and Amp_minPLuc2_Biotin_Rev) corresponding to pMPRAdonor2 (Addgene plasmid #49353) and Kapa HiFi HotStart ReadyMix (Kapa Biosystems). The insert was digested with both KpnI-HF and XbaI (New England Biolabs). Biotinylated primers and undigested inserts were removed as before using Dynabeads M-270 Streptavidin (Thermo Fisher Scientific). Ligation of pJDrcEPP_lib and minP-Luc2 inserts was performed at a 1:3 ratio as before but with T7 DNA ligase (New England Biolabs). Ligation product was cleaned-up and transformed as before, plating similar dilutions but instead growing the remainder transformants overnight in 100x LB + Kanamycin (50 μg/mL) (VWR) at 30°C. The next day, dilutions were counted, estimating 15,877,000 and 17,845,000 original transformants for the placement and spacing library and the number and affinity library, respectively. Library vectors with insert (pJDrcEPP_lib_minPLuc2) were isolated from the remainder transformants using Qiagen Plasmid Plus Maxi Kit (Qiagen).

### Barcode mapping

Barcodes were associated with each library member by sequencing amplicons isolated from pJDrcEPP_lib. 0.5 ng of plasmid was amplified using primers with P5 and P7 Illumina flow cell adapter sequences (Libseq_P7_For and Libseq_P5_Rev) and KAPA HiFi HotStart Real-time PCR Master Mix (2X) (Kapa Biosystems) for 17 cycles. Amplicons were isolated on a 2% TAE agarose gel and bands were confirmed using Agilent’s D1000 ScreenTape and reagents (Agilent Technologies) on a 2200 Tapestation system. Libraries were sequenced on an Illumina MiSeq with a v3 600-cycle reagent kit (Illumina, San Diego, CA) using custom read 1 primer LibSeq_R1Seq_Rev and custom read 2 primer LibSeq_R2Seq_For loaded into the cartridges read 1 and read 2 primer wells, respectively. 35,355,712 reads passed filter and 31,486,576 reads were merged with BBMerge version 9.00. A custom python script was used to map unique barcodes to variants lacking synthesis errors. Briefly, this script searched the last 150 bp of merged reads for sequences perfectly matching the variants designed. The first 20 bp of each merged read was determined to be a barcode and each barcode was then mapped to the most common sequence associated with it, only retaining barcodes that appeared more than twice in merged reads. In order to differentiate between mapped barcodes that are associated with variants with sequencing errors and another variant in the library, we used a Levenshtein distance cut-off of 13 between variants that share a common barcode. This cut-off represented 1% of the total bootstrapped distances between perfect variants in the library. Barcodes mapped to perfect variants were kept if all other variants associated with a barcode fell below this cut-off, retaining 724668 barcodes.

### Library integration

2.6 × 10^6^ Hek293T H11 landing pad cells were plated per T75 flask, 6 flasks in total, and grown in DMEM with 1% Penicillin-streptomycin and 10% FBS (Thermo Fisher Scientific); this is the cell media used in all tissue culture work unless otherwise stated. The next day, the cells were transfected with a total of 187.5 μL Lipofectamine 3000 (Thermo Fisher Scientific), 6.252 mL Opti-MEM (Thermo Fisher Scientific), 13.86 μg BxB1 expression vector (Duportet et al., 2014), 153.36 μg spacing and distance library vector, 180 μg number and affinity library vector and 250 μL P3000 (Thermo Fisher Scientific) following the recommended protocol. BxB1 was added to the DNA mixture at a 1:8 ratio, while both libraries were added at 3x to increase efficiency. Cells were passaged after 3 days onto a T875, in which the media was changed to 1 μg/mL puromycin (Life Technologies, Carlsbad, CA) selection media. Unless otherwise stated, cells were passaged in all tissue culture work according to: trypsinization with Trypsin-EDTA 0.25% (Thermo Fisher Scientific) followed by inactivation with 2x volume of cell media, pelleting at 1000 × g for 5 minutes and resuspension in fresh media. 1/160 of the cells were removed before selection and grown without puromycin to analyze overall integration efficiency. The selection cells were passaged at 1:10, 1:20 or at 1:1 as needed during selection every 1-4 days, with 6 passages in total over 16 days of selection. Cells plated for integration efficiency analysis were passaged at 1:10 or 1:20 every 3 or 4 days for a total of 6 passages. Both cells were analyzed using flow cytometry 20 days after transfection, shown in Supplemental Figure 1B. Samples were prepared in PBS pH 7.4 (Thermo Fisher Scientific) using the LSRII at the UCLA Eli & Edythe Broad Center of Regenerative Medicine & Stem Cell Research Flow Cytometry Core. Cytometer settings were adjusted to: FSC – 157 V, SSC – 233 V, Alexa Fluor 488 – 400V. Selected cells were frozen at 5 × 10^6^ cells/mL in 5% DMSO (Thermo Fisher Scientific) and aliquots were used in the genomic MPRA.

### Luminescence assays

For the landing pad orientation luminescence assay (Supplemental Fig. 1C), 22 × 10^3^ cells containing integrated control sequences were plated in triplicate across a 96-well plate. 100x forskolin stocks were made via serial dilution in DMSO (Thermo Fisher Scientific), and 1x forskolin solutions were made in CD 293 media (Thermo Fisher Scientific) supplemented with 4 mM L-Glutamine (Thermo Fisher Scientific). The next day, media was removed from all 96-wells and replaced with 25 μL of media with forskolin (0, 0.5, 1, 5, 10, 50, 100, and 120 μM). After 4 hours, fluorescence was measured using the Dual-Glo Luciferase Assay Kit (Promega, Madison, WI), in which 10 μL of Dual-Glo Luciferase Reagent was added, cells were shook for 10 minutes and luminescence was measured on a plate reader.

For the MPRA library luminescence assays (Supplemental Fig. 1D), 880,000 H11 landing pad cells and the genomic MPRA cells were resuspended in 12 mL media and 100 μL was distributed per well across a 96-well plate. The next day, the H11 landing pad cells were transfected with a total of 6.6 μL Lipofectamine 3000, 220 μL Opti-MEM, 0.44 μg Renilla luciferase expression vector, 2.15 μg spacing and distance library vector, 2.79 μg number and affinity library vector and 8.8 μL P3000 (Thermo Fisher Scientific) following the recommended protocol, using 10 μL of this mixture per 96-well. 100x forskolin stocks were made via serial dilution in DMSO (Thermo Fisher Scientific), and 1x forskolin solutions were made in cell media. Media was removed from all 96-wells and replaced with 25 μL of media with forskolin (0, 1, 2, 4, 8, 16, and 25 μM). After 4 hours, fluorescence was measured using the Dual-Glo Luciferase Assay Kit (Promega), in which 10 μL of Dual-Glo Luciferase Reagent was added, cells were shook for 10 minutes and luminescence was measured on a plate reader. Renilla luminescence from the transfected cells was measured following Stop & Glo Reagent addition.

### Episomal MPRA

Two mL of a 1.026 × 10^5^ cells/mL stock of H11 Landing Pad cells was plated per 6-well per biological replicate. The next day, cells were transfected with a total of 64.5 μL Lipofectamine 3000, 4.3 mL Opti-MEM, 18.1 μg CRE Spacing and Distance library vector, 21.9 μg CRE Number and Affinity library vector and 86 μL P3000 (Thermo Fisher Scientific) following the recommended protocol, using 250 μL of this mixture per 6-well. Both library vectors were concentrated using a Wizard SV Gel and PCR Clean-up (Thermo Fisher Scientific) prior to transfection. 100x forskolin stocks were made via serial dilution in DMSO (Thermo Fisher Scientific), and 2x forskolin solutions were made in cell media. The next day, 2 mL of media with 2x forskolin was added to the 2 mL media within each 6-well (final concentrations: 0, 2^−5^, 2^−4^, 2^−3^, 2^−2^, 2^−1^, 2^0^ and 2^2^ μM forskolin). After 3 hours, RNA was collected using Qiagen RNeasy Mini Kits with Qiashredder and on-column DNase I digestions with Qiagen RNase-free DNase Set (Qiagen).

Per sample, RNA was reverse-transcribed using 1.5x the recommended materials for Superscript IV Reverse Transcriptase (Thermo Fisher Scientific) with 7.5 μg total RNA and the library-specific primer Creb_Hand_RT, which anneals downstream of barcoded transcripts. The recommended protocol was followed with changes including reverse transcription at 55°C for 1 hour and RNase H (Thermo Fisher Scientific) removal of RNA in RNA:DNA hybrids. To ensure the same amount of barcoded cDNA was used per PCR across forskolin concentrations and that this amount covered library complexity, a preliminary qPCR of total RNA samples was performed alongside serial dilutions of a known amount of barcoded cDNA previously amplified. Sample Cq’s were referenced to Cq’s of the serial dilutions to determine approximate concentrations of barcoded transcripts per total RNA loaded. Volumes of samples that approximated 6000-fold coverage of the number of variants (not including barcode complexity) were determined and used in the following PCR.

cDNA was amplified with NEBNext Q5 Hot Start HiFi PCR Master Mix (New England Biolabs) using input amounts determined from qPCR, distributed across 4 replicates each so as to not exceed 10% of the total PCR volume, and with primers specific to luciferase (Creb_Seq_Luc_R) and a 20 nt annealing site added during reverse-transcription (Creb_Hand). PCR conditions were followed as recommended with 61°C annealing for 20s, extension at 72°C for 20s and a final extension at 72°C for 2 minutes for a total of 10 cycles. Meanwhile, library plasmids mixed at the same ratio as for transfection were used for DNA normalization in the episomal MPRA. 128 ng of this mixture was used per PCR with NEBNext Q5 Hot Start HiFi PCR Master Mix (New England Biolabs) along with the reverse-transcriptase primer (Creb_Hand_RT) and the primer specific to luciferase (Creb_Seq_Luc_R) using the same PCR conditions for cDNA for 9 cycles. Amplicons were isolated on a 2% TAE gel, after which they were cleaned-up again.

A second PCR was performed on both the cDNA and plasmid DNA amplicons in order to add P5 and P7 Illumina flow cell adapter sequences and indices. NEBNext Q5 Hot Start HiFi PCR Master Mix (New England Biolabs) was used with 0.5 ng input and the primers P5_Seq_Luc_F and P7_Ind_#_Han or P7_In_####_Han, with # corresponding to the index code per sample indicated in Supplemental Table 2. PCR conditions were followed as recommended with 63°C annealing for 20s, extension at 72°C for 25s and a final extension at 72°C for 2 minutes for a total of 7 cycles. Bands were confirmed using Agilent’s D1000 ScreenTape and reagents (Agilent Technologies) on a 2200 Tapestation system. Samples were mixed equally and sequenced on a NextSeq500 at the UCLA Technology Center for Genomics & Bioinformatics (TCGB) using the 1 × 75 v2 kit (Illumina) with only 30 cycles. Before sequencing, the custom read 1 sequencing primer Creb_R1_Seq_P and indexing primer Creb_Ind_Seq_P were loaded into the read 1 and index primer positions in the NextSeq cartridge. 365.67 × 10^6^ reads passing filter were obtained across the 8 dilutions (2 replicates each) and plasmid DNA sample with reads per index ranging between 12 × 10^6^ and 20 × 10^6^.

The episomal MPRA performed at concentrations beyond 1 μM forskolin indicated in Supplemental Fig. 2 was similarly transfected and prepped according to the above protocol but with incubations at 0, 2^0^, 2^1^, 2^2^, 2^3^, 2^4^, 25, 2^5^ and 2^6^ μM forskolin. Samples were sequenced on the NextSeq500 using the 1 × 75 v2 kit (Illumina) with 75 cycles and 305.06 × 10^6^ reads passing filter were obtained across the 9 dilutions (2 replicates each) and plasmid DNA sample with reads per index ranging between 10 × 10^6^ and 14 × 10^6^. The single CRE library indicated in Supplemental Figure 3 was similarly transfected and prepped according to the above protocol but with incubations at 0 and 25 μM forskolin. Additionally, volumes of cDNA samples that approximated 1500-fold coverage of the number of variants (not including barcode complexity) were determined and used in the following PCR. 101.55 × 10^6^ HiSeq reads passing filter were obtained across all cDNA and plasmid DNA samples.

### Genomic MPRA

2.5 × 10^5^ integrated and selected cells were plated on 2 separate 6-wells, forming the two biological replicates used in the MPRA. These cells were passaged twice in their expansion, after which cells were frozen at 5 × 10^6^ cells/mL in 5% DMSO (Thermo Fisher Scientific). For the MPRA, 5 aliquots of each replicate, 2.5 × 10^7^ cells total, were thawed and grown to cover the initial bottleneck amount 100-fold. 2 days later these cells were trypsinized and plated for stimulation at 3.47 × 10^6 cells per 150 cm plate with 20 mL of media, eight plates total per replicate. Two days later, both replicates were stimulated by adding 20 mL of media with 16 μM forskolin to the 20 mL media already on plates. After 3 hours, replicates were trypsinized, combined, spun down at 1000 × g for 5 minutes, resuspended in media and split evenly into 2 tubes, one RNA extraction and one for genomic DNA extraction.

Cells aliquoted for RNA processing were spun down and resuspended in 3.2 mL of RLT (1% ß-Mercaptoethanol) from a Qiagen RNeasy Midi kit (Qiagen). Cells were homogenized by passing the lysate through an 18-gauge needle 10 times and were stored at −80°C. Two days later, lysates were thawed and processed according to the RNeasy Midi Protocol for Isolation of Total RNA from Animal Cells from Qiagen RNeasy Midi/Maxi Handbook (09/2010) (Qiagen) starting at the addition of 1x volume of 70% ethanol to thawed lysates. On-column DNase I digestions were performed with the Qiagen RNase-free DNase Set (Qiagen). RNA was eluted with 200 μL RNAse-free water (Qiagen) and subsequently concentrated using an Amicon Ultra-0.5 mL Centrifugal Filter with a 10 kDa cut-off. Total RNA was stored at −20°C.

Cells aliquoted for genomic DNA were spun down and resuspended in PBS pH 7.4 (Thermo Fisher Scientific) twice to give a final concentration of 1 × 10^7^ cells/mL. Samples were processed according to the Sample Preparation and Lysis Protocol for Cell Cultures from QIAGEN Genomic DNA Handbook (08/2001) using the settings for the Qiagen Blood and Cell Culture DNA Maxi Kit (Qiagen). Pelleted nuclei were frozen at −20°C before G2 buffer was added. Two days later, nuclei were thawed and the remainder of the protocol was followed using Qiagen Protease digestion at 50°C for 60 minutes, and precipitating DNA according to the recommended protocol for vortexing and centrifugation after isopropanol addition followed by washing with cold 70% ethanol. Genomic DNA was resuspended in 800 μL Qiagen Elution Buffer (Qiagen) and left at room temperature overnight. The next day, gDNA was shook at 600 rpm for 3 hours at 55°C. RNase A (DNase and protease-free, Thermo Fisher Scientific) was added to a final concentration of 99 ng/μL. Over the next 3 days, 600 μL Qiagen Elution Buffer (Qiagen) was added incrementally, with additional shaking at 55-60°C after each addition for a total of 28 hours; this was largely due to resuspension issus. Resuspended DNA was stored at 4°C.

Per replicate, 130 μg total RNA was reverse transcribed using the recommended materials for Superscript IV Reverse Transcriptase (Thermo Fisher Scientific) but with 10 μg total RNA instead of the recommended 5 μg per 20 μL reaction. The library-specific primer Creb_RT_Hand_3, which anneals downstream of barcoded transcripts, was added to reactions and the recommended protocol was followed with changes including reverse transcription at 55°C for 1 hour and RNase H (Thermo Fisher Scientific) removal of RNA in RNA:DNA hybrids. RNAse A (DNase and protease-free, Thermo Fisher Scientific) was added to each reaction at a final concentration of 100 ng/μL and incubated at 37°C for 30 minutes. Reactions were combined and concentrated using an Amicon Ultra-0.5 mL Centrifugal Filter with a 10 kDa cut-off (Sigma-Aldrich). To ensure the same amount of barcoded cDNA was used per PCR across replicates and that this amount covered library complexity, a preliminary qPCR of total RNA samples was performed alongside serial dilutions of a known amount of barcoded cDNA previously amplified. Sample Cq’s were referenced to Cq’s of the serial dilutions to determine approximate concentrations of barcoded transcripts per total RNA loaded per replicate. Of the 30 μL volume of cDNA remaining in each replicate, total barcoded molecules were estimated. 30 μL of the replicate with the lower concentration of barcoded molecules was used in the following PCR while a portion of the replicate with the higher concentration was used to approximate similar cDNA input into the following PCR. Both amounts loaded covered the original 2.5 × 10^5^ cell bottleneck amount 24.5-fold.

cDNA was amplified with NEBNext Q5 Hot Start HiFi PCR Master Mix (New England Biolabs) using volumes determined from qPCR distributed across 5 replicates each, so as not to exceed 10% of the total PCR volume, and with primers specific to luciferase (Creb_Luc_Seq_R) and a 20 nt annealing site added during reverse-transcription (Creb_Hand). PCR conditions were followed as recommended with 61°C annealing for 20s, extension at 72°C for 20s and a final extension at 72°C for 2 minutes for a total of 16 cycles. A second PCR was performed in order to add P5 and P7 illumina flow cell adapter sequences and indices. NEBNext Q5 Hot Start HiFi PCR Master Mix (New England Biolabs) was used with 0.5 ng cDNA and the primers P5_Seq_Luc_F and P7_Ind_##_Han, with ## corresponding to the index code per sample indicated in Supplemental Table 2. PCR conditions were followed as recommended with 63°C annealing for 20s, extension at 72°C for 25s and a final extension at 72°C for 2 minutes for a total of 7 cycles.

Meanwhile, gDNA was aliquoted into 2 tubes evenly before PCR to establish 2 technical replicates per biological replicate. Per technical replicate, gDNA was amplified with NEBNext Q5 Hot Start HiFi PCR Master Mix (New England Biolabs) with a biotinylated reverse-transcriptase primer Creb_Hand_RT_3 and a biotinylated primer specific to luciferase (Creb_Luc_Seq_R). 5 μg gDNA was loaded per 50 μL in a 96-well PCR plate, with 57 total reactions per technical replicate for one biological replicate and only 51 for the other, due to sample loss. PCR conditions were followed as recommended with 61°C annealing for 20s, extension at 72°C for 20s and a final extension at 72°C for 2 minutes. After 7 cycles, wells were combined and cleaned-up using Agencourt AMPure XP beads (Beckman Coulter, Brea, CA). 0.4x volume of beads were used per sample and after magnetic separation, the supernatant was collected, isolating the amplicons from genomic DNA. Similar as in (Matreyek et al., 2017), 40% of eluted volume was used in a second PCR to add P5 and P7 illumina flow cell adapter sequences and indices. This volume was distributed across 14 replicates per technical replicate so as to not exceed 10% of the total PCR volume and amplified using NEBNext Q5 Hot Start HiFi PCR Master Mix (New England Biolabs) with the primers P5_Seq_Luc_F and P7_Ind_##_Han, with ## corresponding to the index code per sample. PCR conditions were followed as recommended with 63°C annealing for 20s, extension at 72°C for 25s and a final extension at 72°C for 2 minutes for a total of 14 cycles.

Both the cDNA and gDNA amplicons were isolated on a 2% TAE gel and bands were confirmed using Agilent’s D1000 ScreenTape and reagents (Agilent) on a 2200 Tapestation system. Samples were mixed equally and sequenced on a HiSeq2500 at UCLA’s Broad Stem Cell Research Center using a 1 × 50 kit. Before sequencing, the custom read 1 sequencing primer Creb_R1_Seq_P and indexing primer Creb_Ind_Seq_P were loaded into the read 1 and index primer positions in the HiSeq cartridge. Indexed samples were de-multiplexed in-house using a custom python script that matched indices with those submitted and all 1 bp mutations, accounting for sequencing errors. 143,759,096 reads passing filter were obtained across the 2 biological replicates, each consisting of 2 genomic DNA technical replicates and 1 RNA sample. Reads per index ranged between 9 × 10^6^ − 22 × 10^6^.

### Processing MPRA Sequencing Data

A custom python script was used to isolate barcode sequences from reads and determine their total number of reads. Briefly, the first 20 sequences were extracted from each read, reverse complemented to match the barcode format from barcode mapping and then the occurence of each of these barcodes was summed as their total number of reads per indexed sample. RStudio (R version 3.5.3 and the packages: tidyverse 1.2.1, lemon 0.4.3, viridisLite 0.3.0, cowplot 0.9.4, caTools 1.17.1.2, broom, 0.5.1, and modelr 0.1.4) was used for the remainder of data processing. Barcodes were normalized to sequencing depth per sample and represented as normalized reads per million. Barcodes were retained and used in variant expression determination only if they also appeared in the barcode-variant mapping table. Barcodes that were present in the barcode-variant mapping table that were not present in a sample were given the value of 0 normalized reads per million amongst retained barcodes.

### Determining MPRA Variant Expression

In the episomal MPRA, barcodes were retained across all samples with >6 reads in the DNA sample. Barcode expression was determined by dividing normalized barcode reads per million in RNA samples to their normalized reads in the plasmid DNA sample. Variants were retained across all samples if they had >7 barcodes retained in the plasmid DNA sample. Median expression per variant was determined by taking the median expression of all barcodes associated with a single variant, maintaining variants with >0 expression in all samples. In total, 4162 of the original 4185 variants designed along with the CRE control were retained after processing. Median variant expression in the single CRE library indicated in Supplemental Fig. 3 was similarly determined except retaining barcodes across all samples with > 5 reads in the DNA sample.

In the genomic MPRA, expression was calculated similarly as for the episomal MPRA, with changes accounting for 2 DNA technical replicates. Barcodes were retained across all samples per biological replicate if they had > 6 reads in both DNA technical replicates. Barcode expression was determined by dividing normalized barcode reads per million in RNA samples to their average normalized reads across the genomic DNA samples. Per biological replicate, variants were similarly retained in the RNA sample if they were associated with >7 barcodes retained in the combined DNA sample. Median expression per variant was determined by taking the median expression of all barcodes associated with a single variant. Variants were retained for further analysis if they had >0 expression in both biological replicates. Overall, 3479 of the original 4185 variants designed were retained in analysis in addition to the CRE control. Variants not retained consisted of: 128 from the number and affinity library and 578 from the spacing and distance library (488 of this was with background 41, of which was dropped from analysis). For both MPRAs, the average expression between biological replicates was used for all variant analyses.

### CREB∷ßzip structure superpositions along DNA

The structure coordinates of a Creb∷ßzip dimer bound to the somatostatin CRE (Schumacher et al., 2000) was downloaded from PDB (code: 1DH3) and loaded into Coot (version 0.8.9.2). Models of CREB protein spacing were made by using the LSQ Superpose function, superpositioning a copy of the protein:DNA complex onto the original structure using least squares fit to the mainchain of Chain B, corresponding to one DNA strand. Briefly, a model of 5 bp CREB protein spacing was established by taking residues −10:−4 on Chain B of the reference structure and moving them to residues 4:10 on Chain B. This matched 63 atoms with a rms deviation of 1.22. A model of 10 bp CREB protein spacing was established by taking residues −11:-9 on Chain B of the reference structure and moving them to residues 8:10 on Chain B. This matched 26 atoms with an rms deviation of 1.49.

A model of six CREB proteins bound to six CREs in the Number and Affinity library was constructed similarly. CREB protein bound to the first CRE was created by taking residues 5:9 on Chain B of the reference structure and moving them to residues −9:5 on Chain B. CREB protein bound to the second CRE was created by taking residues −9:5 on Chain B of the reference structure and moving them to residues 4:8 on Chain B. The third instance of CREB protein-CRE was created by taking residues −9:−5 on Chain B of the reference structure of the second CREB protein-CRE and moving them to residues 5:9 on Chain B then taking residues −9:5 on Chain B of this new reference structure and moving them to residues 4:8 on Chain B. This was repeated sequentially using the previous CREB protein-CRE structure as a reference until superpositioning the sixth instance of CREB protein bound to a CRE. A total of 10 LSQ superpositions were performed matching 45 atoms each time with a rms deviation of either 1.15 or 1.09 depending upon the reference and moving residues.

### Log-linear expression modelling

A model was fit using `lm()` in R *stats* package to predict average expression from the independent contribution of background and the 6 CRE positions in the site number and affinity library (expression ~ background + site1 + site2 + site3 + site4 + site5 + site6). Background and each position was represented as categorical variables according to the 3 backgrounds used and 3 possible affinities per position (consensus CRE, weak CRE, and no CRE). The percent of variance explained per model term was obtained using the sum of squares from `anova()`.

## Supplemental Materials

**Supplemental Figure 1.**
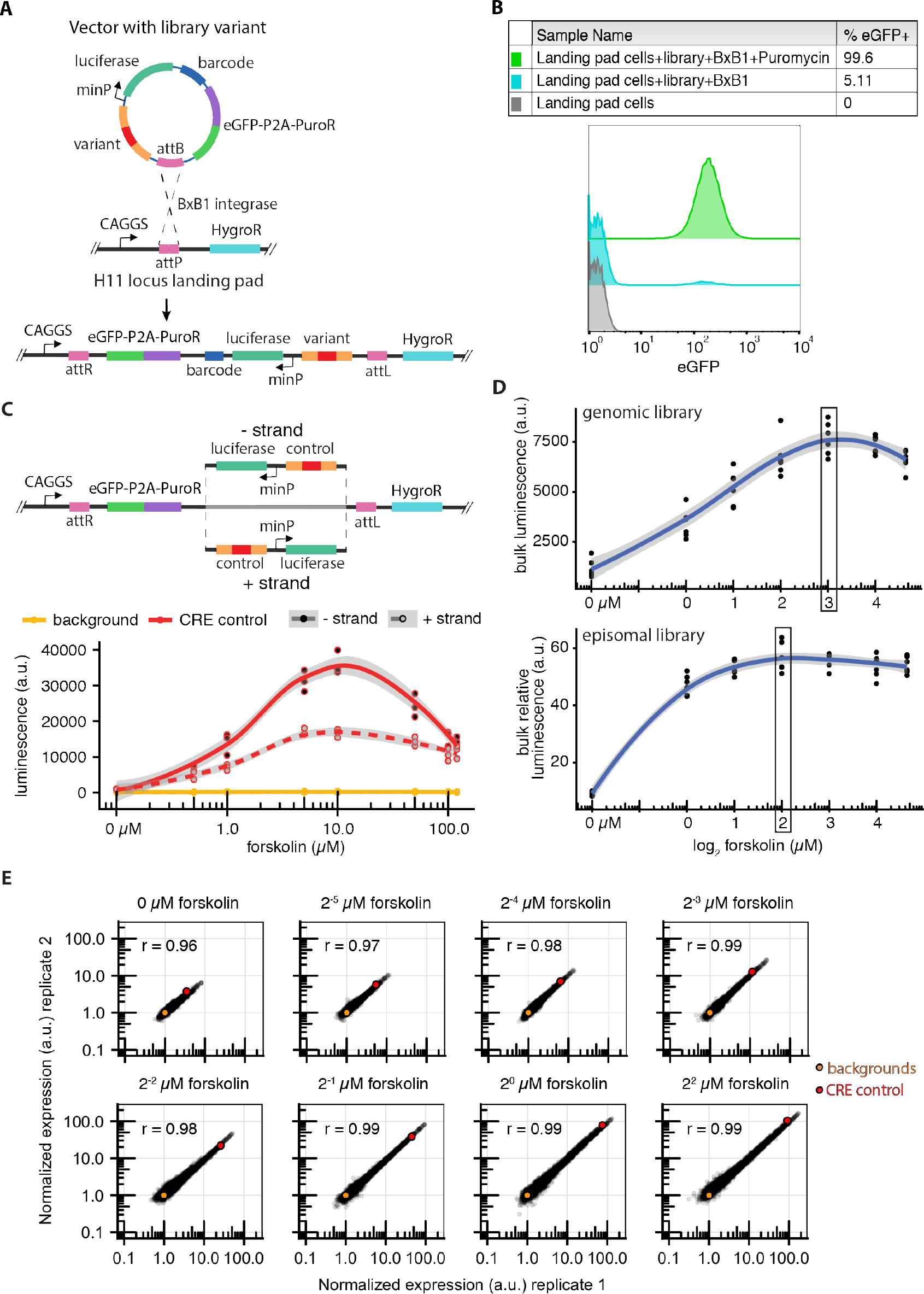
Establishment of the episomal and genomic MPRA. (A) Library-containing vectors are genomically-integrated by co-transfecting with a BxB1 expression vector into a HEK293T cell line containing a single-copy landing pad at the H11 locus. BxB1-mediated integration occurs through the genomic recombination site, attP, and the vector recombination site, attB. Successful integration at H11 switches cell antibiotic resistance from hygromycin to puromycin, via the genomic CAGGS promoter, in addition to driving expression of eGFP. Vector and landing pad components not shown to scale. (B) Integration of library-containing vectors in the genomic MPRA was monitored using eGFP activation upon genomic integration. 5.11% of transfected cells expressed eGFP from an integrated construct (cyan). Successful integrants were isolated after outgrowth in media containing puromycin (green). (C) Integration orientation of library controls at the landing pad resulted in different levels of induced expression. The negative strand orientation placed the attL sequence immediately upstream of the CRE control while in the positive strand orientation, this was replaced by bacterial backbone. The negative strand orientation was chosen for the genomic CRE MPRA. Lines indicate a loess fit with shaded regions indicated standard error. (D, top graph) Bulk genomically-integrated library luciferase expression measured across forskolin dilutions, 6 technical replicates each. The genomic MPRA was performed using 8 μM forskolin for comparisons to the episomal MPRA. (D, bottom graph) Similar bulk luciferase expression measurements but after transfection of the episomal library. Luciferase luminescence normalized to luminescence from a Renilla transfection control. The episomal MPRA analysis was performed at 4 μM forskolin for comparisons to the genomic MPRA. In both graphs, lines indicate a loess fit with shaded regions indicated standard error. (E) Replicability plots following the episomal MPRA titration curve in figure 1D. Expression from each variant was normalized to the expression of their corresponding background per biological replicate to visualize induction following increasing forskolin. Replicability ranged from r = 0.96 to r = 0.99 across concentrations.

**Supplemental Figure 2.**
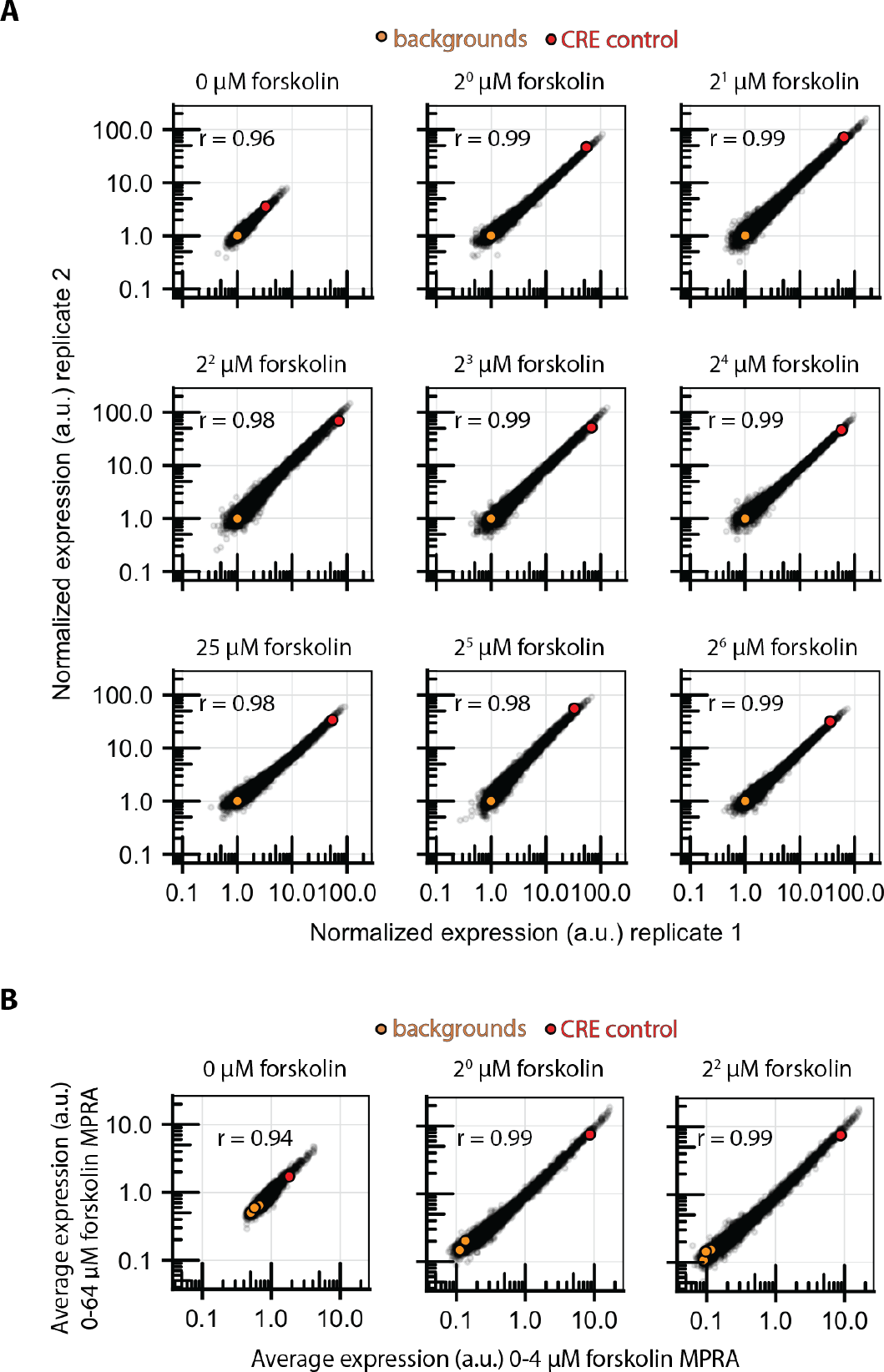
Episomal MPRA tested at concentrations beyond maximal induction indicate little change in expression range. (A) The episomal MPRA was performed at concentrations spanning beyond those used in the main analysis (Figure 1D and Supplemental Fig. 1E) to confirm the full induction range in episomal conditions. (B) Repeated concentrations between episomal MPRAs indicate high reproducibility (r = 0.94-0.99).

**Supplemental Figure 3.**
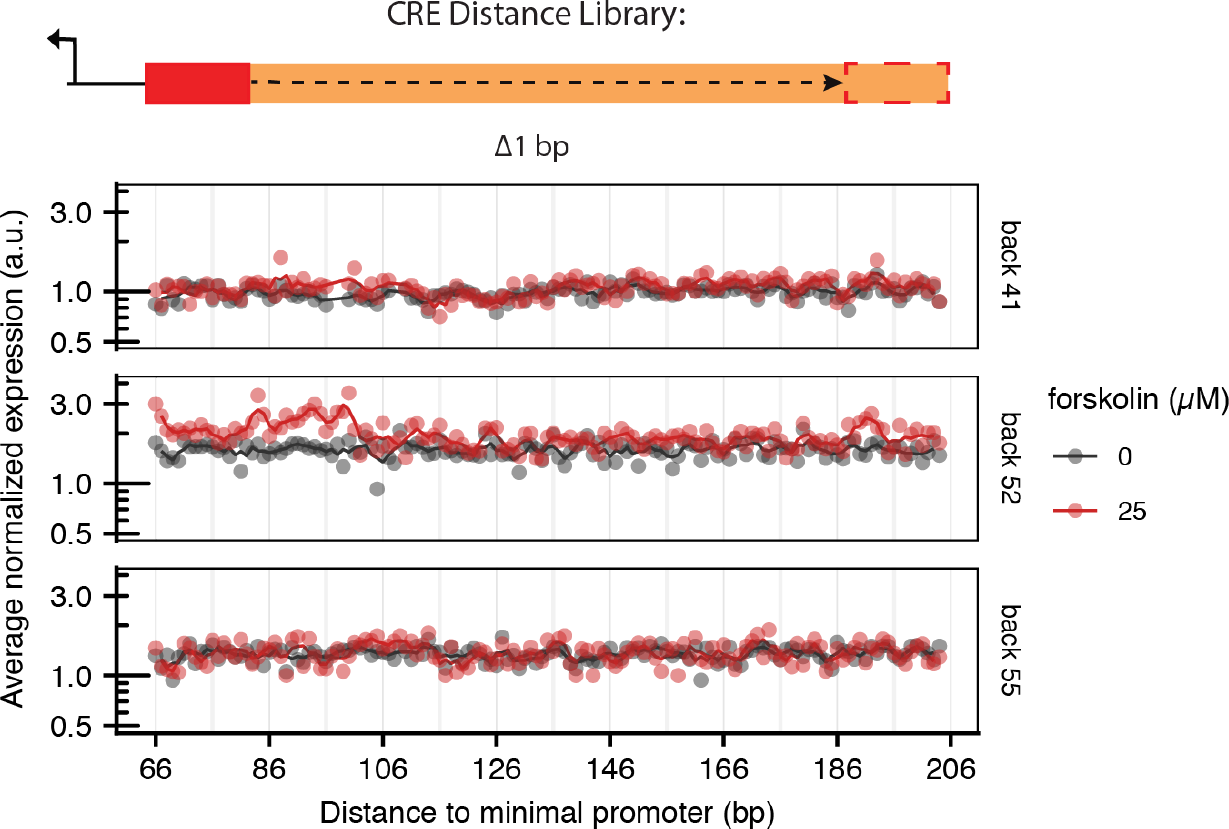
Variants with one CRE drive minimally-induced expression and do not indicate large expression variation as CRE distance to the promoter is varied in the episomal MPRA. Variants containing a single CRE at every position along the three backgrounds assayed in an separate MPRA at uninduced and fully-induced forskolin concentrations. Variant expression is normalized to the expression of their backgrounds, averaged across replicates and compared between forskolin concentrations. Line shown is the 3 bp moving average expression estimate. As the distance of one CRE to the promoter varies there is little expression variation in all but a portion of distances in variants with background 52.

**Supplemental Figure 4.**
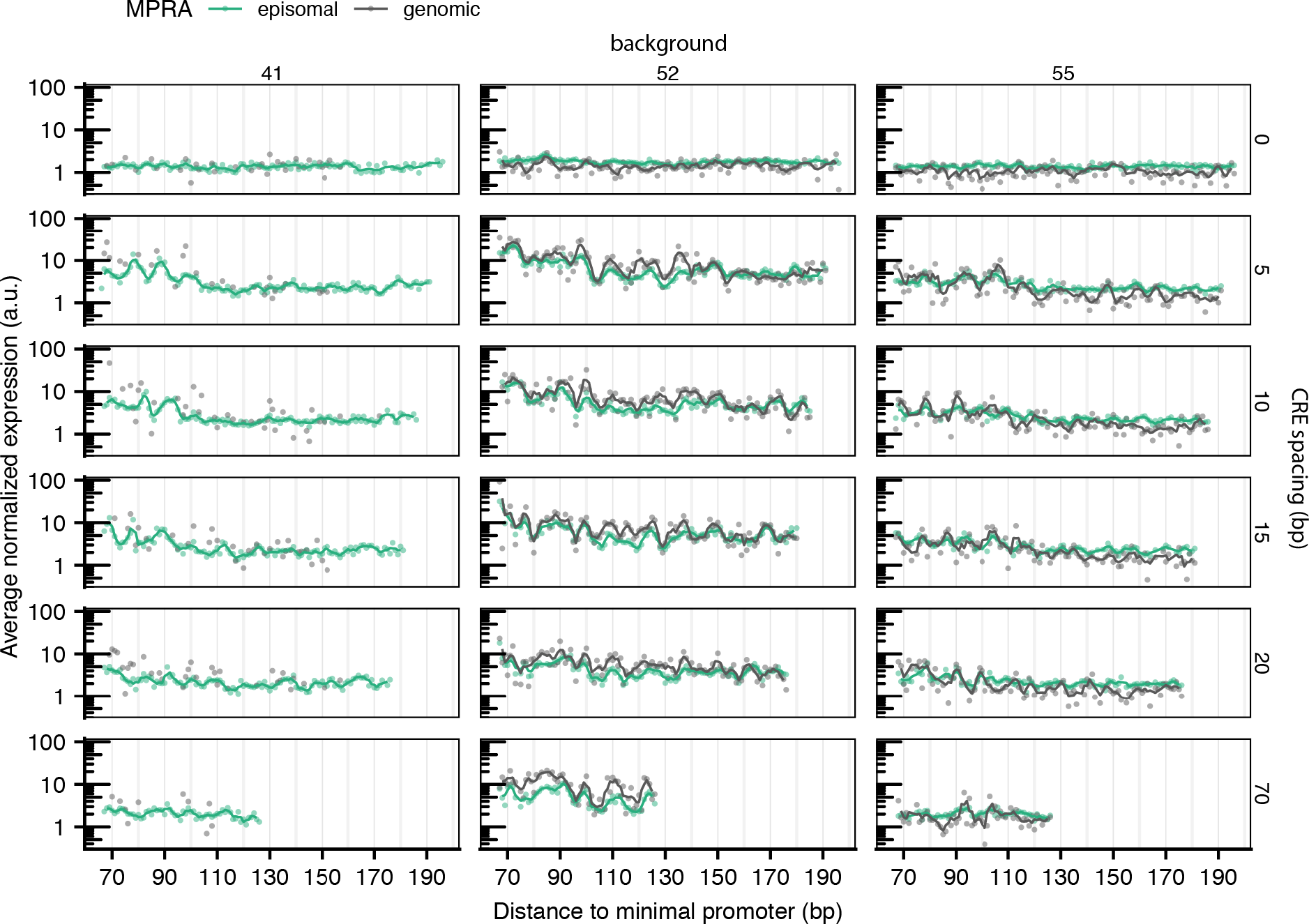
Expression measurements for all variants in the spacing and distance library retained in analysis. All MPRA expression measurements used for analysis of variants in the spacing and distance library. Comparisons to the episomal MPRA were performed with expression obtained at 4 μM forskolin. Data quality filters in the genomic MPRA remove 74% of variants with background 41, thus only episomal MPRA expression is used for periodicity analysis in variants with this background. Similar normalized expression profiles are observed in between variants in the episomal versus the genomic context. Variants with 0 bp CRE spacing were predicted to occlude the binding of CREB protein to both CREs; here we observe minimal expression driven by these variants.

**Supplemental Figure 5.**
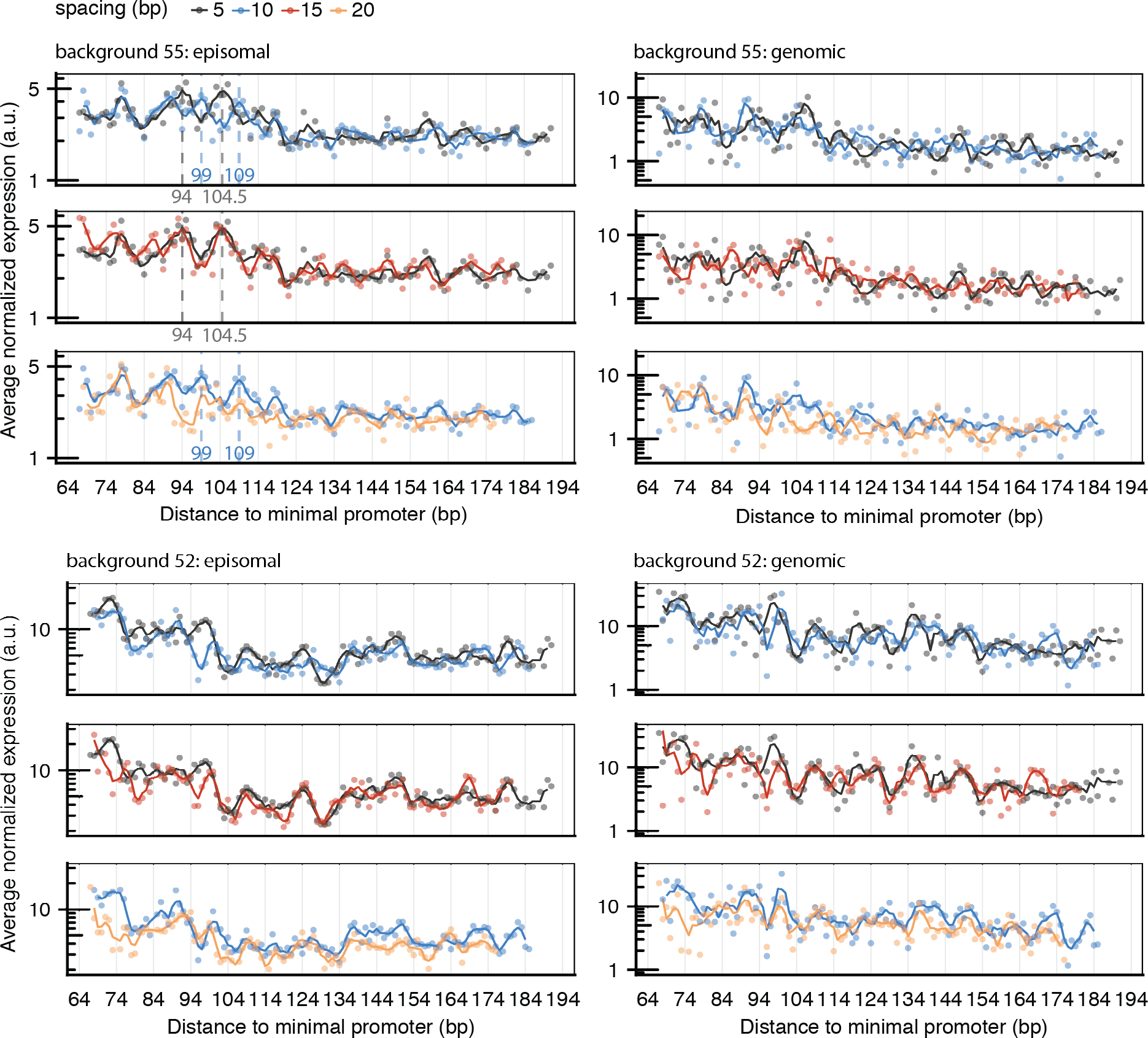
Variant periodicity offset graphs as in Fig. 3A with backgrounds 55 and 52 in both MPRAs. Average variant expression across replicates according to proximal CRE distance to the promoter (x-axis) is subset according to background and MPRA and colored by the spacing between CREs. The line corresponds to a 3 bp moving average estimate. Variants with background 55 in the episomal MPRA display a similar offset between 5 and 10 bp spacings and alignment between both 5 and 15 bp and 10 and 20 bp spacings. Dashed lines correspond points at local expression maxima across CRE distances or the midpoint between points if a maxima was not apparent in variants with 5 and 10 bp CRE spacings. Similar periodicity offsets and alignments are not as pronounced along this background in the genomic MPRA and along background 52 in both MPRAs.

**Supplemental Figure 6.**
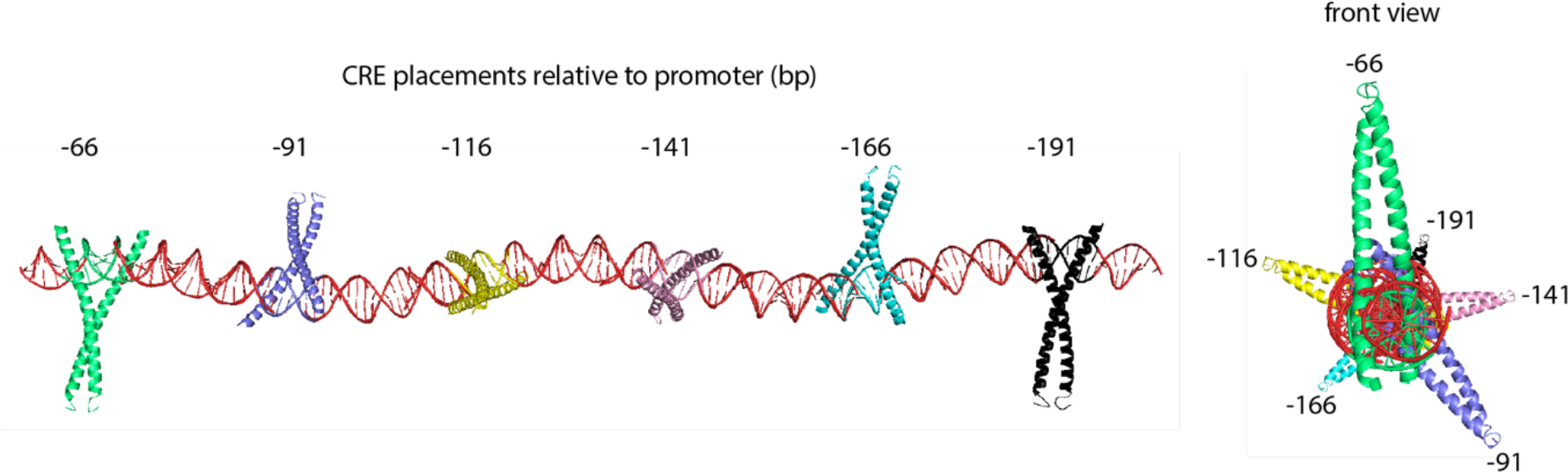
CREs in the Number and Affinity library were placed at positions which were expected to sample multiple CREB protein orientations along DNA. Using the published structure of a CREB∷bZIP dimer bound to CRE (PDB: 1DH3), we modeled the expected placement of CREB dimers bound to the 6 CRE sites and constant 17 bp CRE *spacing* used in the CRE Number and Affinity library. Orientation of each CREB dimer is indicated relative to the first dimer following CRE *distances* relative to the minimal promoter.

**Supplemental Figure 7.**
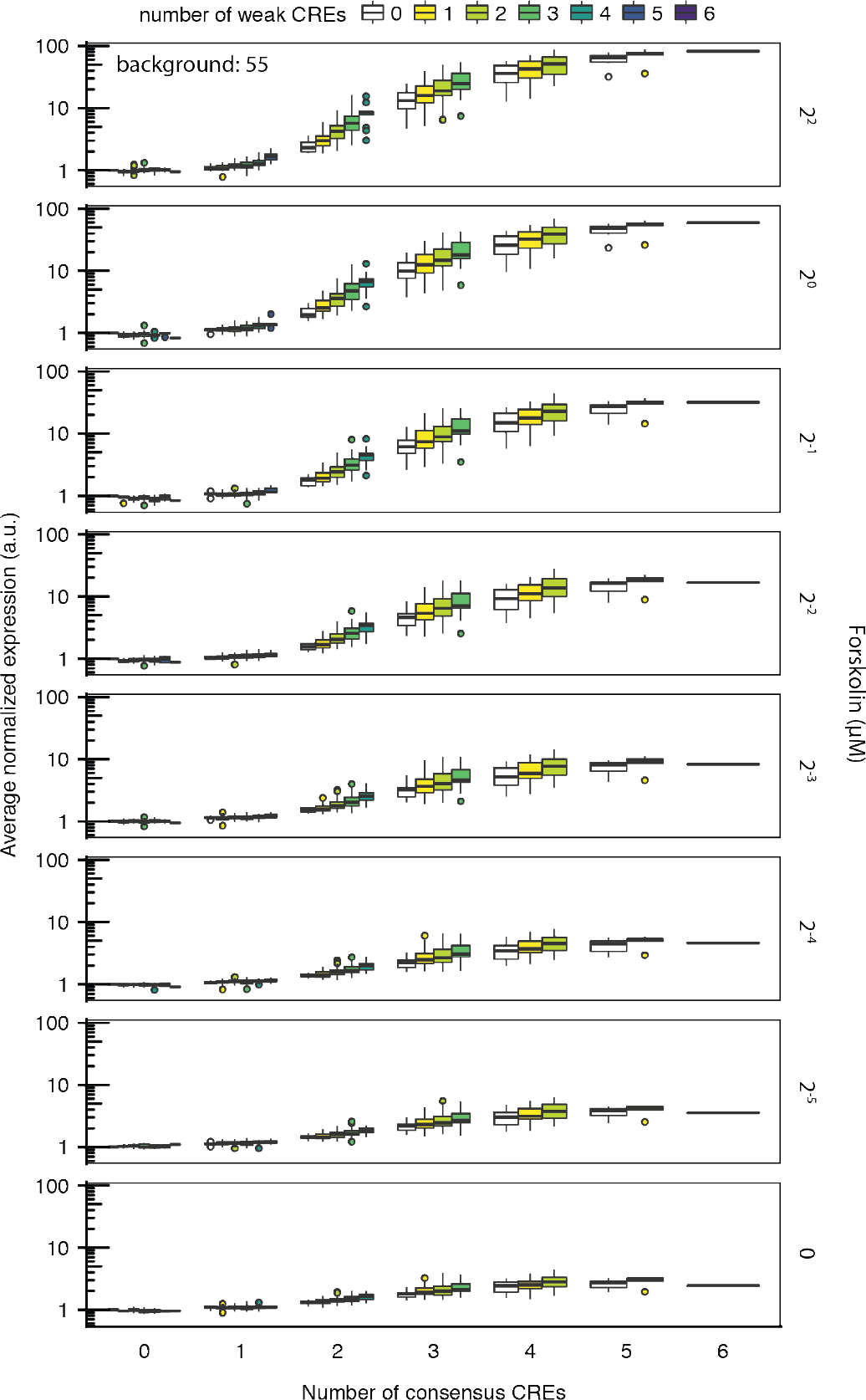
The extent of CREB activation following CRE numbers per variant is determined by CREB activation levels. The episomal expression of variants with background 55 according to their number of consensus (x-axis) and weak (colored subsets) CREs. The extent of this response is modulated by CREB protein activation via forskolin induction. In the absence of forskolin, CRE number drives a modest non-linear increase in expression. Variants with background 41 and 52 followed similar effects across forskolin concentrations (Not shown).

**Supplemental Figure 8.**
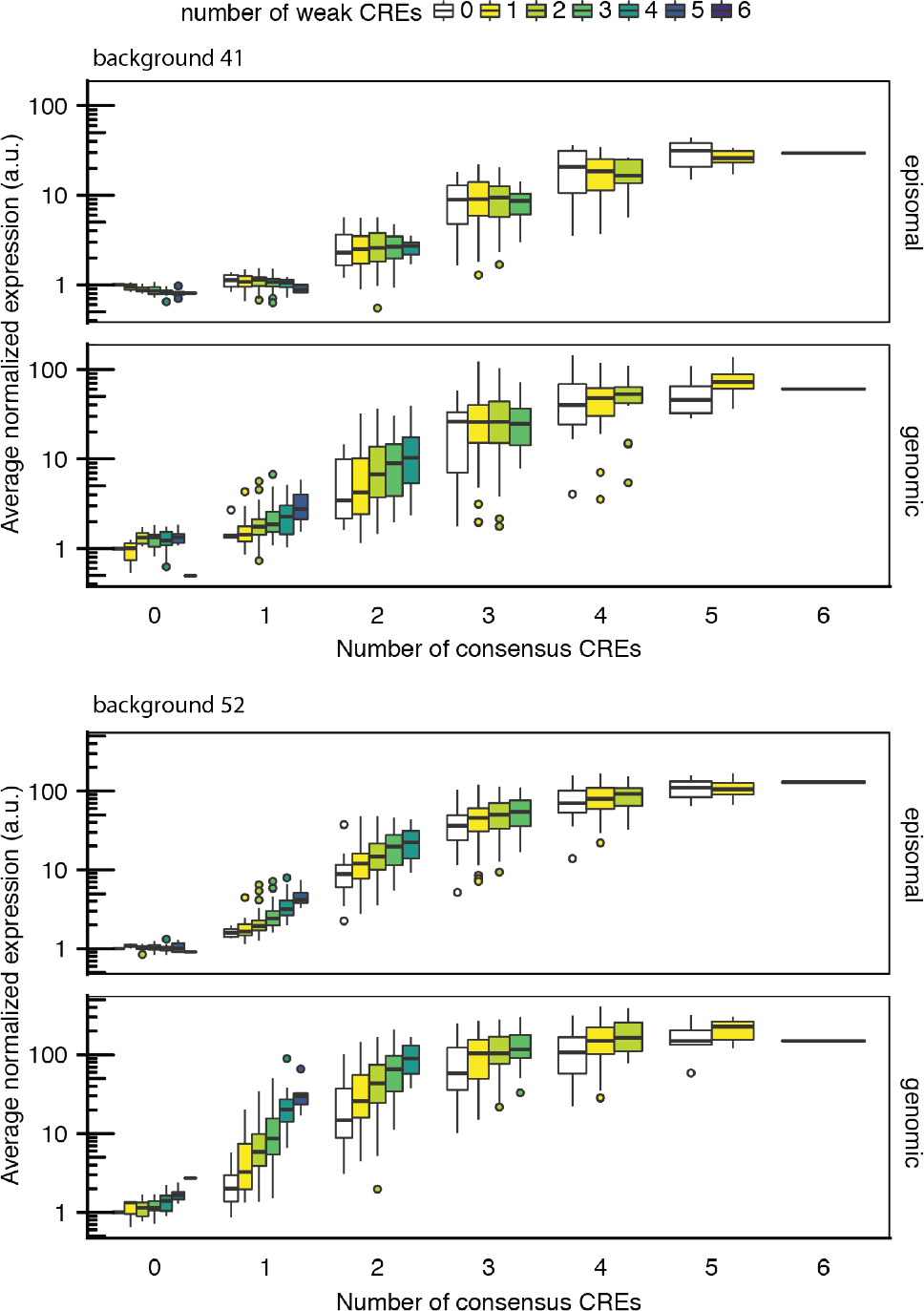
The number of consensus CREs in variants largely determines expression while the number of weak CREs has a variable effect between episomal and genomic MPRAs. Similar as in Fig. 4A, variants with background 41 and 52 in both MPRAs grouped according to their total number of both consensus (x-axis) and weak CREs (colored subsets) and average expression plotted per variant per MPRA (y-axis). The number of consensus CREs largely determines the expression per variant and drives a non-linear trend in expression. For reasons that are not apparent, the number of weak CREs per variant drives a similarly non-linear increase in expression across all variants but those with background 41 in the episomal MPRA.

**Supplemental Figure 9.**
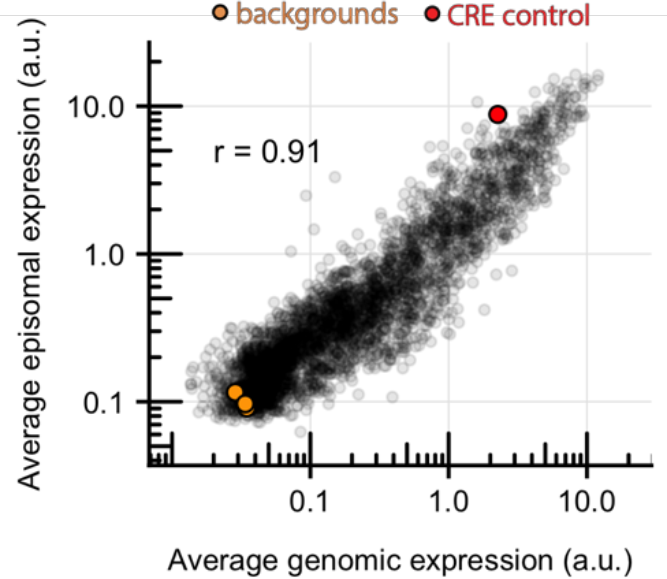
Library member expression largely correlates between MPRA formats. Despite differences, library variant expression correlates well between the episomal and genomic MPRA (r = 0.91).

**Supplemental Table 1.**
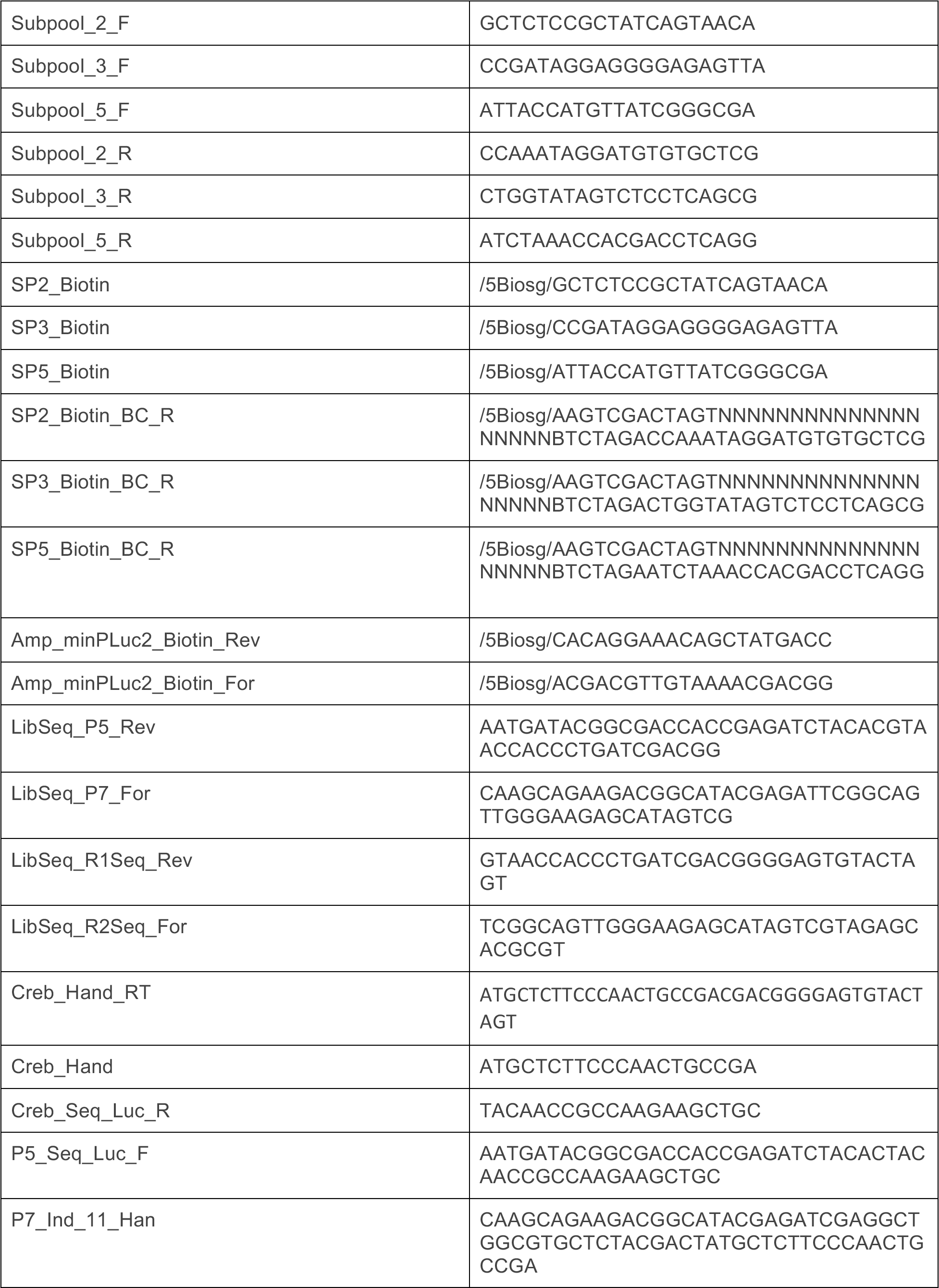
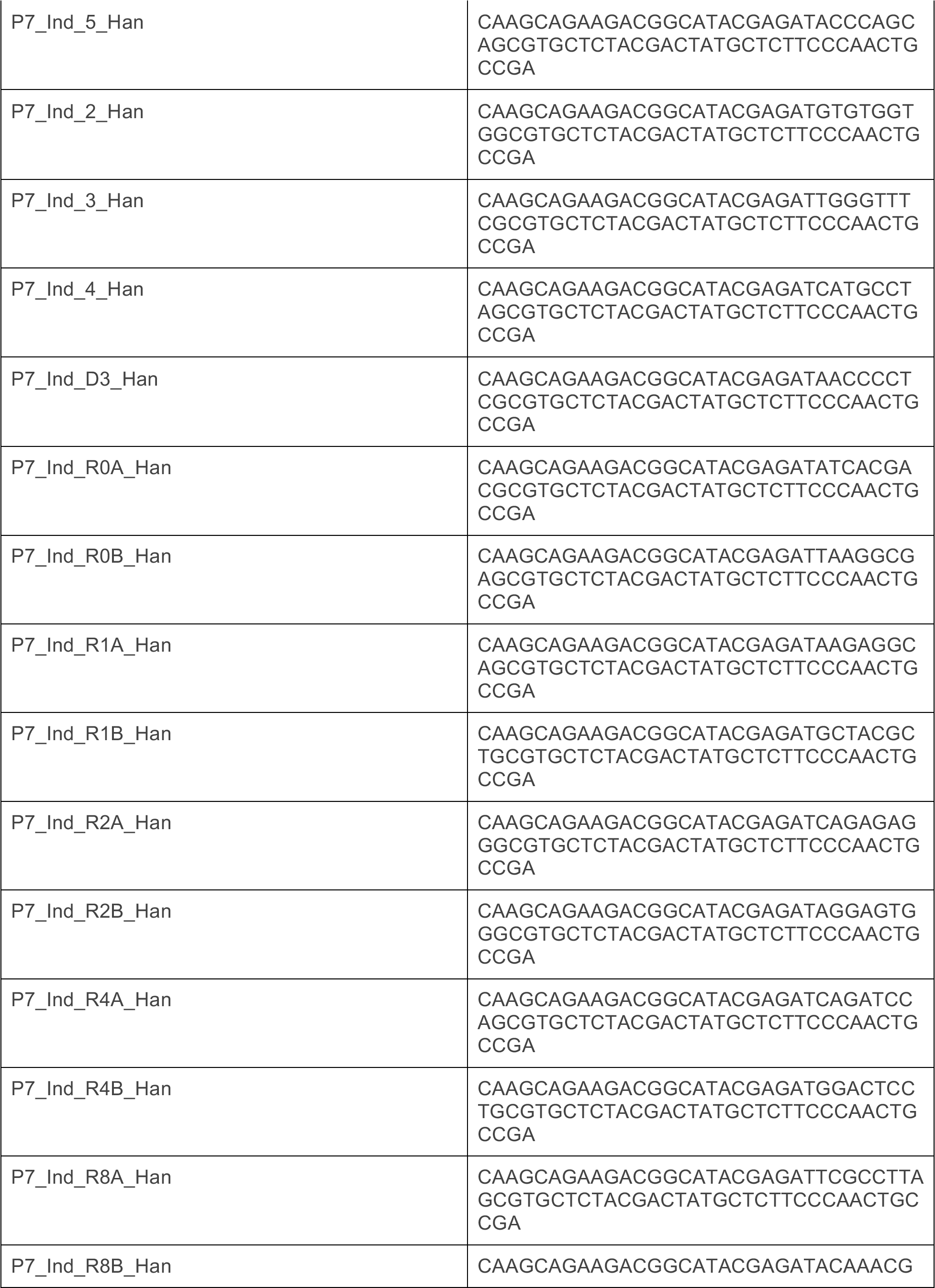
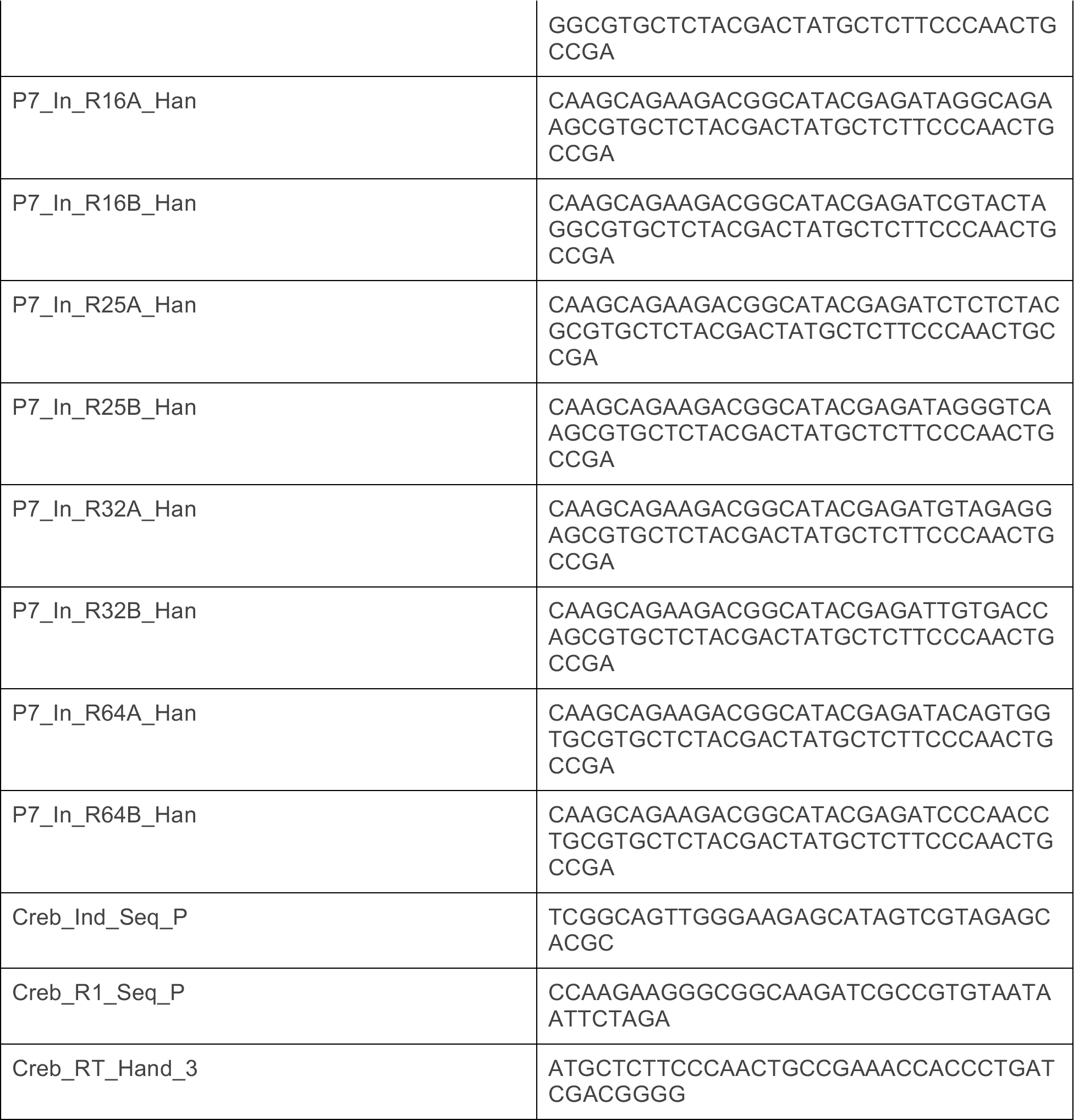
List of primers and sequences used throughout this study.

